# LKB1 inactivation modulates chromatin accessibility to drive metastatic progression

**DOI:** 10.1101/2021.03.29.437560

**Authors:** Sarah E. Pierce, Jeffrey M. Granja, M. Ryan Corces, Jennifer J. Brady, Min K. Tsai, Aubrey B. Pierce, Rui Tang, Pauline Chu, David M. Feldser, Howard Y. Chang, Michael C. Bassik, William J. Greenleaf, Monte M. Winslow

**Author notes:** Correspondence to: S.E.P., W.J.G., or M.M.W. These authors contributed equally to this work (Sarah E. Pierce, Jeffrey M. Granja). **Contact Information:** William J. Greenleaf, PhD, Stanford University School of Medicine, 279 Campus Drive, Beckman Center B257, Stanford, CA 94305-5168, Monte M. Winslow, PhD, Stanford University School of Medicine, 279 Campus Drive, Beckman Center B256, Stanford, CA 94305-5168.

## Abstract

Metastasis is the leading cause of cancer-related deaths, enabling cancer cells to expand to secondary sites and compromise organ function^1^. Given that primary tumors and metastases often share the same constellation of driver mutations^2–4^, the mechanisms driving their distinct phenotypes are unclear. Here, we show that inactivation of the frequently mutated tumor suppressor gene, liver kinase B1 (LKB1), has evolving effects throughout lung cancer progression, leading to the differential epigenetic re-programming of early-stage primary tumors compared to late-stage metastases. By integrating genome-scale CRISPR/Cas9 screening with bulk and single-cell multi-omic analyses, we unexpectedly identify LKB1 as a master regulator of chromatin accessibility in lung adenocarcinoma primary tumors. Using an in vivo model of metastatic progression, we further reveal that loss of LKB1 activates the early endoderm transcription factor SOX17 in metastases and a metastatic-like sub-population of cancer cells within primary tumors. SOX17 expression is necessary and sufficient to drive a second wave of epigenetic changes in LKB1-deficient cells that enhances metastatic ability. Overall, our study demonstrates how the downstream effects of an individual driver mutation can appear to change throughout cancer development, with implications for stage-specific therapeutic resistance mechanisms and the gene regulatory underpinnings of metastatic evolution.

## Main

The serine/threonine kinase LKB1 (also known as STK11) is frequently inactivated in many cancer types, including pancreatic, ovarian, and lung carcinomas, and germline heterozygous LKB1 mutations cause Peutz-Jeghers familial cancer syndrome^5–7^. Loss of LKB1 leads to both increased primary tumor growth and the acquisition of metastatic ability in lung adenocarcinoma, the most common subtype of lung cancer^8–12^. However, beyond its well-established role as an activator of AMPK-related kinases^10,13–16^, the mechanisms by which LKB1 constrains metastatic ability and cell state are unclear.

In addition to exhibiting high rates of LKB1 mutations, human lung adenocarcinomas frequently harbor mutations in chromatin modifying genes, such as *SETD2*, *ARID1A*, and *SMARCA4*^5^, suggesting that genetic alterations can drive tumor progression by influencing epigenetic state. This interplay between genetic and epigenetic mechanisms is starting to be characterized in other cancer types; for example, H3K27M mutations in diffuse midline gliomas suppress epigenetic repressive capacity and differentiation^17^ and SDH-deficiency in gastrointestinal stromal tumors initiates global DNA hyper-methylation and unique oncogenic programs^18^. However, the role of chromatin dynamics in regulating lung adenocarcinoma progression, and particularly metastatic spread, remains uncharacterized.

### Generation of LKB1-deficient lung adenocarcinoma cell lines with restorable alleles of *Lkb1*

To establish a tractable platform to assess LKB1 function in cancer, we generated cell lines from oncogenic KRAS-driven, TP53-deficient (*Kras^G12D^;p53^−/−^; KP*) murine lung tumors harboring homozygous restorable alleles of *Lkb1* (*Lkb1^TR/TR^*) and a tamoxifen-inducible FlpOER allele (*Rosa26^FlpOER^*) (**Extended Data Figs. 1a, b**; see Methods)^19^. In LKB1-restorable cell lines (LR1 and LR2), a gene trap cassette within intron 1 of *Lkb1* introduces a splice acceptor site and premature transcription-termination signal before any sequences encoding functional domains of LKB1. Treatment with 4-hydroxytamoxifen (4-OHT) results in FlpOER nuclear translocation and excision of the FRT-flanked gene trap cassette, thereby restoring full-length expression of *Lkb1* (**Extended Data Figs. 1c-e**). Restoring LKB1 decreased proliferation in cells in culture and decreased tumor growth after transplantation into mice, whereas treating LKB1-unrestorable,FlpOER-negative cell lines (LU1 and LU2) with 4-OHT had no effect (**Extended Data Figs 1f-i**).

### Identification of a genetic link between the LKB1/SIK pathway and chromatin regulation

To identify genes and pathways that contribute to LKB1-mediated tumor suppression, we performed a proliferation-based genome-scale CRISPR/Cas9 knock-out screen in both LKB1-deficient and LKB1-restored cells (**Fig. 1a**). We first transduced a Cas9-expressing LKB1-restorable cell line (LR1;Cas9) with a lentiviral library containing ~10 sgRNAs per gene in the genome as well as ~13,000 inert controls^20^. After selecting for transduced cells, we treated the cells with 4-OHT or vehicle for 12 days and sequenced the sgRNA region of the integrated lentiviral vectors (**Supplementary Table 1 and Extended Data Fig. 1j**). As expected, the most highly enriched sgRNA target in LKB1-restored cells compared to LKB1-deficient cells was *Lkb1* itself (**Fig. 1b and Extended Data Fig. 1k**). Gene Ontology (GO) term enrichment analysis^21^ of the remaining top targets surprisingly revealed a strong enrichment of chromatin-related processes (**Fig. 1c**). In particular, six of the top 20 targets were chromatin modifiers (*Suv39h1, Arid1a, Eed, Suz12, Trim28, and Smarce1*) (**Fig. 1b and Extended Data Fig. 1l**), suggesting that the LKB1 pathway engages chromatin regulatory mechanisms to limit growth in lung cancer.

**Figure 1.**
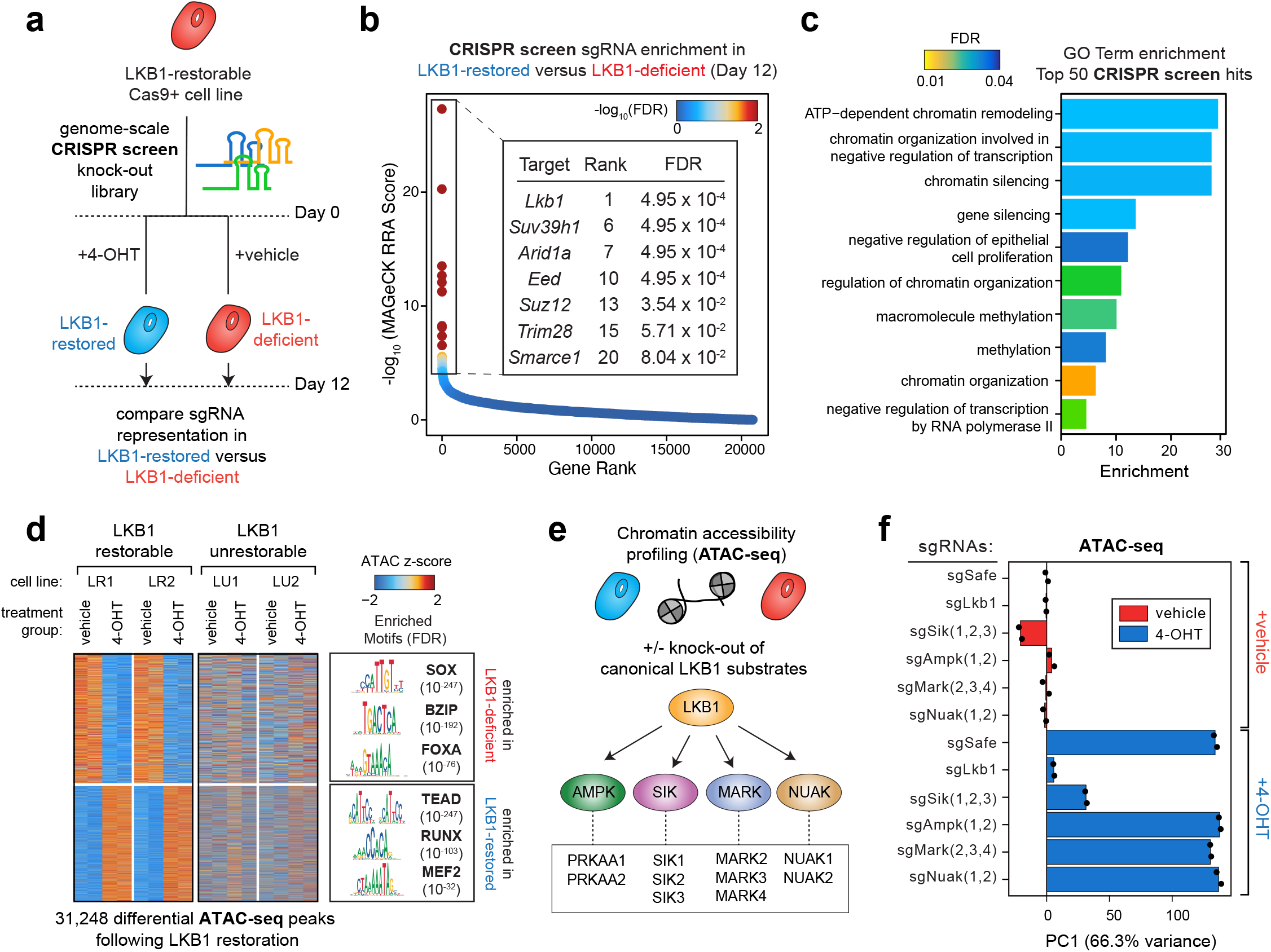
An LKB1-SIK axis regulates chromatin accessibility in lung adenocarcinoma. **a.** Schematic of a genome-scale screen in an LKB1-restorable, Cas9+ lung adenocarcinoma cell line (LR1;Cas9) treated with 4-OHT to restore LKB1 or treated with vehicle to remain LKB1-deficient (see also Extended Data Fig. 1a). **b.** sgRNA targets (genes) enriched in LKB1-restored versus LKB1-deficient cells are ranked by log_10_(MAGeCK RRA score) and colored by −log_10_ FDR values. LKB1 and six chromatin-related genes with an FDR < 0.01 are shown alongside their individual rank and FDR values. **c.** PANTHER GO term enrichment of the top 50 sgRNA targets (genes) enriched in LKB1-restored cells compared to LKB1-deficient cells. Bars are colored by enrichment FDR values. **d.** *Left*: Heatmap of chromatin peak accessibility for each cell line after treatment with 4-OHT or vehicle for six days. Each row represents a z-score of log_2_ normalized accessibility within each cell line using ATAC-seq. *Right*: Transcription factor hypergeometric motif enrichment with FDR indicated in parentheses. **e.** Schematic of knocking out canonical LKB1 substrate families with arrays of sgRNAs in an LKB1-restorable cell line (LR1;Cas9), treating with 4-OHT or vehicle for six days, and performing ATAC-seq. **f.** Principle component analysis (PCA) of the top 10,000 variable ATAC-seq peaks across the twelve indicated sgRNA populations that were treated with either 4-OHT or vehicle for six days. Individual principle components besides PC1 (66.3%) account for <4% of the variance in the dataset.

To understand how LKB1 expression affects chromatin accessibility, we performed the assay for transposase-accessible chromatin using sequencing (ATAC-seq) on LKB1-deficient and LKB1-restored cells from two cell lines (LR1;Cas9 and LR2;Cas9) (**Fig. 1d, Extended Data Fig.2a-c,** and **Supplementary Table 2**)^22,23^. Remarkably, LKB1 restoration resulted in consistent, large-scale chromatin accessibility changes, with >14,000 regions increasing and >16,000 regions decreasing in accessibility (**Fig. 1d** and **Extended Data Fig. 2d**). LKB1-induced chromatin changes were of similar magnitude to the overarching chromatin accessibility differences between cancer sub-types, such as basal and luminal breast cancer (**Extended Data Fig. 2e**)^24,25^. In addition, the majority of LKB1-induced chromatin changes occurred within 24-48 hours of LKB1 restoration (**Extended Data Fig. 2f-j**), suggesting rapid regulation by the LKB1 pathway. Genomic regions with increased accessibility in LKB1-restored cells were enriched for TEAD and RUNX transcription factor binding motifs, whereas genomic regions with increased accessibility in LKB1-deficient cells were enriched for SOX and FOXA motifs (**Fig. 1d** and **Extended Data Fig. 2h**). Interestingly, inactivating the top chromatin modifier hits from the screen (*Eed*, *Suz12*, *Trim28*, *Suv39h1*) in the LR1;Cas9 cell line appears to delay, but not prevent, LKB1-induced chromatin accessibility changes (**Extended Data Fig. 3**), suggesting compensation between chromatin regulatory pathways.

The canonical tumor suppressive role for LKB1 involves the phosphorylation and activation of AMPK-related kinases, including the AMPK, SIK, NUAK, and MARK families^13^. To evaluate whether the downstream substrates of LKB1 contribute to LKB1-induced chromatin changes, we knocked out each family with multiple arrays of sgRNAs and performed ATAC-seq with and without LKB1 restoration in the LR1;Cas9 cell line (**Fig. 1e and Extended Data Fig.4a**). Knocking out the *Sik* family (*Sik1*, *Sik2*, and *Sik3* simultaneously) almost entirely abrogated the ability of LKB1 to induce chromatin accessibility changes (**Fig. 1f and Extended Data Figs.4b-g**), whereas inactivation of the *Ampk*, *Nuak*, or *Mark* families or the individual *Sik* paralogs (*Sik1*, *Sik2*, or *Sik3* independently) had no effect (**Fig. 1f and Extended Data Figs. 4b-j**). Therefore, the SIK family of kinases act redundantly, but collectively mediate LKB1-induced chromatin changes.

### LKB1 mutation status defines chromatin accessibility sub-types of human lung adenocarcinoma

Given the strength of LKB1/SIK-induced chromatin accessibility changes in the murine restoration model, we next evaluated whether LKB1 mutation status correlates with chromatin accessibility differences across human lung adenocarcinoma primary tumors. De novo hierarchical clustering of the 21 lung adenocarcinoma samples from the TCGA ATAC-seq dataset^25^ revealed two novel chromatin sub-types of lung cancer (annotated as Chromatin Type 1 and Chromatin Type 2) (**Fig. 2a**). Of the top ~200 mutated genes in lung adenocarcinoma, LKB1 was the most significantly enriched mutated gene in Chromatin Type 2 tumors compared to Chromatin Type 1 tumors (FDR = 0.088) (**Extended Data Fig. 5a**).

**Figure 2.**
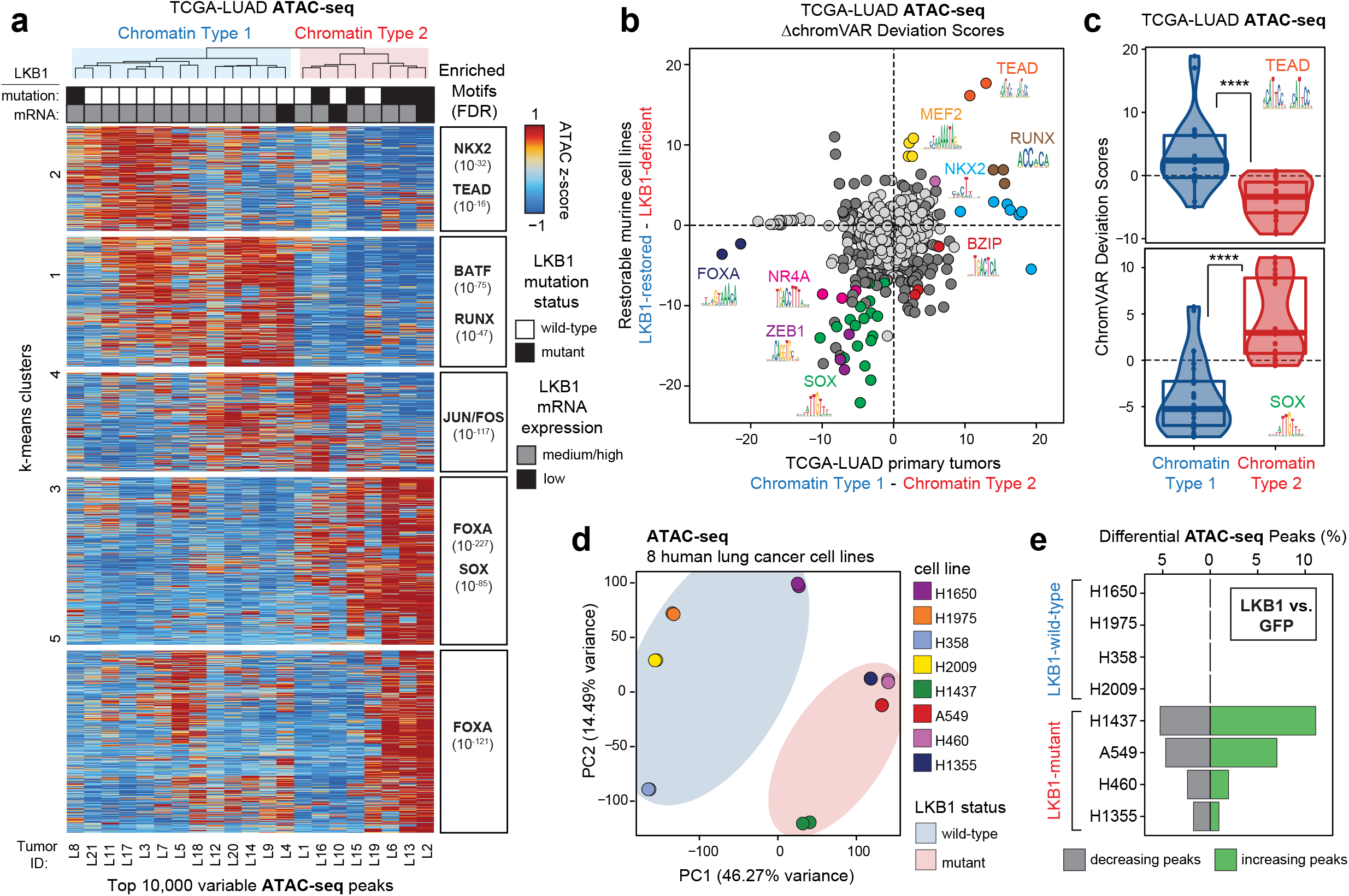
LKB1 mutation status distinguishes the two main chromatin sub-types of human lung adenocarcinoma. **a.** *Left*: Unsupervised hierarchical clustering of 21 human lung adenocarcinoma samples from the TCGA-ATAC dataset using the top 10,000 variable peaks across all samples, visualized as a heatmap of peak accessibility. Each row represents a z-score of log_2_ normalized accessibility. *Right*: Transcription factor hypergeometric motif enrichment in each k-means cluster with FDR indicated in parentheses. **b.** Comparison of the changes in motif accessibility (ΔchromVAR Deviation Scores) across Chromatin Type 1 and Chromatin Type 2 human primary tumors (x-axis) and across LKB1-restored and LKB1-deficient murine cells (x-axis). Dark grey or colored points are significantly different (q < 0.05, see Methods) across both comparisons. Light grey points are not significant. Only motifs for transcription factors shared across human and murine CisBP databases are shown. **c.** ChromVAR deviation scores for TEAD (*top*) and SOX (*bottom*) transcription factor motifs for samples in the TCGA-LUAD ATAC-seq dataset. ****p<10^−6^ using a two-sided t-test. **d.** PCA of the top 10,000 variable ATAC-seq peaks across eight human lung cancer cell lines. LKB1 mutant status is indicated. **e.** Percent of differential ATAC-seq peaks (|log_2_fold change| > 0.5, FDR < 0.05) in cells transduced to express LKB1 compared to a GFP control.

We next evaluated how the distinct chromatin accessibility states of Chromatin Type 1 and Chromatin Type 2 human primary tumors compared to the acute chromatin accessibility changes induced by LKB1 restoration in murine cells. We first calculated the differential accessibility of transcription factor binding motifs between Chromatin Type 1 and Type 2 human tumors and between LKB1-restored and LKB1-deficient murine cells using chromVAR^26^. For motifs that were conserved across murine and human datasets, we then compared their differential motif deviation scores (**Fig. 2b**). Overall the differences between Chromatin Type 1 and Type 2 primary tumors were highly concordant with the differences between LKB1-restored and LKB1-deficient murine lung cancer cells. In particular, genomic regions containing TEAD and RUNX motifs were more accessible in Chromatin Type 1 tumors and LKB1-restored murine cells, and genomic regions containing SOX and FOXA motifs were more accessible in Chromatin Type 2 tumors and LKB1-deficient murine cells (**Fig. 2b, c and Extended Data Fig. 5b**). These results suggest that LKB1 mutations are not only enriched in Chromatin Type 2 tumors, but also that inactivation of LKB1 is likely a defining feature that divides lung adenocarcinoma into two chromatin accessibility sub-types.

To further evaluate LKB1-dependent effects on chromatin, we performed ATAC-seq on a panel of eight human non-small cell lung cancer cell lines (H1650, H1975, H358, H2009, H1437, A549, H460, H1355). Principle component analysis (PCA) and hierarchical clustering unbiasedly stratified LKB1-wild-type and LKB1-mutant cell lines based on their chromatin profiles (**Fig. 2d and Extended Data Fig. 6a**). Genomic regions containing RUNX and TEAD motifs were more accessible in LKB1-wild-type cell lines, whereas genomic regions containing SOX and FOXA motifs were more accessible in LKB1-mutant cell lines (**Extended Data Figs. 6b-c**). Furthermore, similar to the murine restoration model, expressing wild-type LKB1 in LKB1-mutant human lung cancer cells dramatically altered chromatin accessibility, with on average >15,000 regions increasing and >10,000 regions decreasing in accessibility (**Fig. 2e and Extended Data Fig. 6d**). The magnitude of differential accessibility changes was positively correlated with the baseline LKB1-deficiency gene expression score of each cell line (R=0.96) (**Extended Data Fig. 6e**)^27^. Expression of an orthogonal tumor suppressor KEAP1 in KEAP1-mutant cell lines (A549, H460, and H1355) induced very minor chromatin changes (**Extended Data Figs. 6f-h**), emphasizing the specificity of the LKB1 tumor suppressor pathway in regulating chromatin accessibility states in lung cancer.

### LKB1 activity governs chromatin accessibility in murine lung adenocarcinoma primary tumors and metastases

LKB1-deficiency cooperates with oncogenic KRAS in mouse models of lung adenocarcinoma to promote both early-stage tumor growth and late-stage metastasis^8^. To determine whether LKB1 loss has stage-specific effects on tumor progression, we generated an *in vivo* model system to directly compare LKB1-proficient and LKB1-deficient primary tumors and metastases. We incorporated homozygous *Lkb1* floxed alleles into the metastatic, *Kras^LSL-G12D/+^;p53^flox/flox^;Rosa26^LSL-tdTomato^* (*KPT*) mouse model to maintain a common genetic background between LKB1-proficient and LKB1-deficient tumors. Lentiviral Cre administration into the lungs of *KPT* and *KPT;Lkb1^flox/flox^* mice led to the development of aggressive primary tumors capable of seeding spontaneous metastases within 4-7 months. Overall, metastases were observed in 90% of *KPT;Lkb1^−/−^* mice and ~60% of KPT mice (**Supplementary Table 5**). We FACS-isolated cancer cells from individual primary tumors and metastases and performed ATAC-seq (n=12 *KPT* primary tumors, 13 *KPT;Lkb1^−/−^* primary tumors, 4 *KPT* metastases, and 5 *KPT;Lkb1^−/−^* metastases; **Fig. 3a**). PCA of the 25 primary tumors stratified samples based on LKB1 status, similar to the stratification of Chromatin Type 1 and Chromatin Type 2 human primary tumors (**Fig. 3b**). In addition, the motif accessibility differences between LKB1-proficient and LKB1-deficient murine samples were consistent in directionality with our previous datasets (**Extended Data Fig. 7a, b**), with SOX motifs more accessible in LKB1-deficient samples and TEAD, RUNX, and MEF2 binding sites more accessible in LKB1-proficient samples. These results underscore the robustness of LKB1-driven chromatin accessibility states across species and model systems.

**Figure 3.**
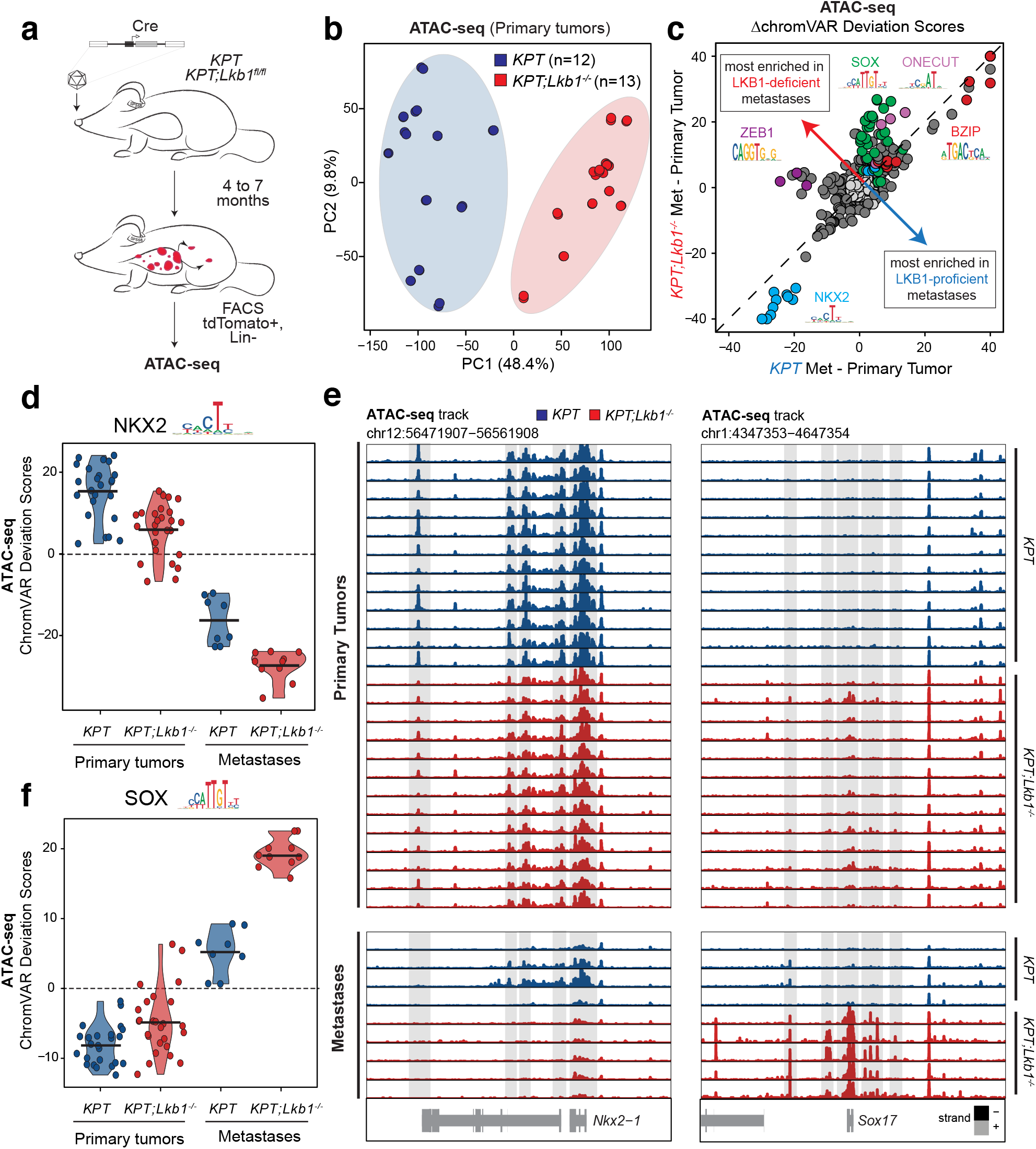
Genotype-specific activation of SOX17 expression in metastatic, LKB1-deficient cells. **a.** Schematic of tumor initiation, sample processing, and multi-omic profiling. Lentiviral Cre initiates tumors in *Kras^LSL-G12D^;p53^flox/flox^;Rosa26^LSL-tdTomato^* (*KPT*) mice with and without homozygous *Lkb1^flox/flox^* alleles. tdTomato+ cancer cells negative for the lineage markers CD45, CD31, F4/F80, and Ter119 were sorted by FACS before library preparation for ATAC-seq, scATAC-seq, and RNA-seq. **b.** PCA of the top 10,000 variable ATAC-seq peaks across 25 primary tumor samples. Technical replicates are averaged. **c.** Comparison of the changes in motif accessibility (ΔchromVAR Deviation Scores) across LKB1-proficient(x-axis) and LKB1-deficient (y-axis) metastases compared to primary tumors of the same genotype. Dark grey or colored points are called significantly different (q < 0.05, see Methods) across both comparisons. Light grey points are not significant. **d.** chromVAR deviation scores for NKX2 motifs across LKB1-proficient (*KPT*) and LKB1-deficient (*KPT;Lkb1^−/−^*) primary tumor and metastasis samples. **e.** *Nkx2.1* and *Sox17* genome accessibility tracks for each primary tumor (*top*) and metastasis (*bottom*) sample. **f.** chromVAR deviation scores for SOX motifs across LKB1-proficient (*KPT*) and LKB1-deficient (*KPT;Lkb1^−/−^*) primary tumor and metastasis samples.

### LKB1-deficient metastases activate expression of the transcription factor SOX17

To evaluate genotype- and metastasis-specific epigenetic features, we compared the chromatin accessibility profiles of LKB1-proficient and LKB1-deficient metastases after correcting for their related primary tumor chromatin accessibility profiles. Downregulation of the transcription factor *Nkx2.1* has previously been shown to increase metastatic ability in lung adenocarcinoma^28^; similarly, all metastases had decreased local accessibility at the *Nkx2.1* locus, decreased *Nkx2.1* mRNA expression, and decreased accessibility of genomic regions containing NKX2 motifs compared to primary tumors (**Fig. 3c-e** and **Extended Data Fig. 7c**). In contrast, the most prominent genotype-specific difference was that LKB1-deficient metastases had high accessibility of genomic regions containing SOX motifs (**Fig. 3f**). Of all the SOX family members, LKB1-deficient metastases specifically expressed high levels of the early endoderm transcription factor *Sox17*, whereas LKB1-deficient primary tumors expressed low levels of *Sox17* and LKB1-proficient primary tumors and metastases did not express *Sox17* (**Extended Data Fig. 7d**). Similarly, the *Sox17* locus was highly accessible in LKB1-deficient metastases, weakly accessible in LKB1-deficient primary tumors, and inaccessible in LKB1-proficient samples (**Fig. 3e**). Thus, high SOX17 expression and increased accessibility of genomic regions containing SOX motifs correlate with metastatic progression in LKB1-deficient lung adenocarcinoma. While SOX17 has not previously been associated with the LKB1 pathway, express SOX17 in mature lung epithelial cells is sufficient to inhibit differentiation and induce hyperplastic clusters of diverse cell types^29^, suggesting that SOX17 can have strong effects on cell state and behavior.

### LKB1-deficient primary tumors harbor sub-populations of metastatic-like, SOX17+ cells

To characterize the heterogeneity and level of SOX17 protein expression in lung adenocarcinoma, we performed SOX17 immunohistochemistry on LKB1-proficient and LKB1-deficient primary tumors and metastases. LKB1-proficient primary tumors and metastases were universally SOX17-negative, while all LKB1-deficient metastases contained SOX17+ cancer cells (**Fig. 4a** and **Extended Data Figs. 8a, b**). In addition, a fraction of LKB1-deficient primary tumors (63/203 tumors) harbored sub-populations of SOX17+ cells, primarily located within invasive acinar structured areas (**Extended Data Figs. 8a, b**). In support of the hypothesis that LKB1 signaling regulates SOX17 expression, we also found that *LKB1* mRNA expression was negatively correlated with *SOX17* mRNA expression in metastatic lung adenocarcinoma cells derived from human tumors^30^ (**Extended Data Fig. 8c**; R= −0.81). In addition, LKB1-deficient Chromatin Type 2 human primary tumors had higher accessibility at the *SOX17* locus compared to Chromatin Type 1 human primary tumors (**Extended Data Fig. 8d**).

**Figure 4.**
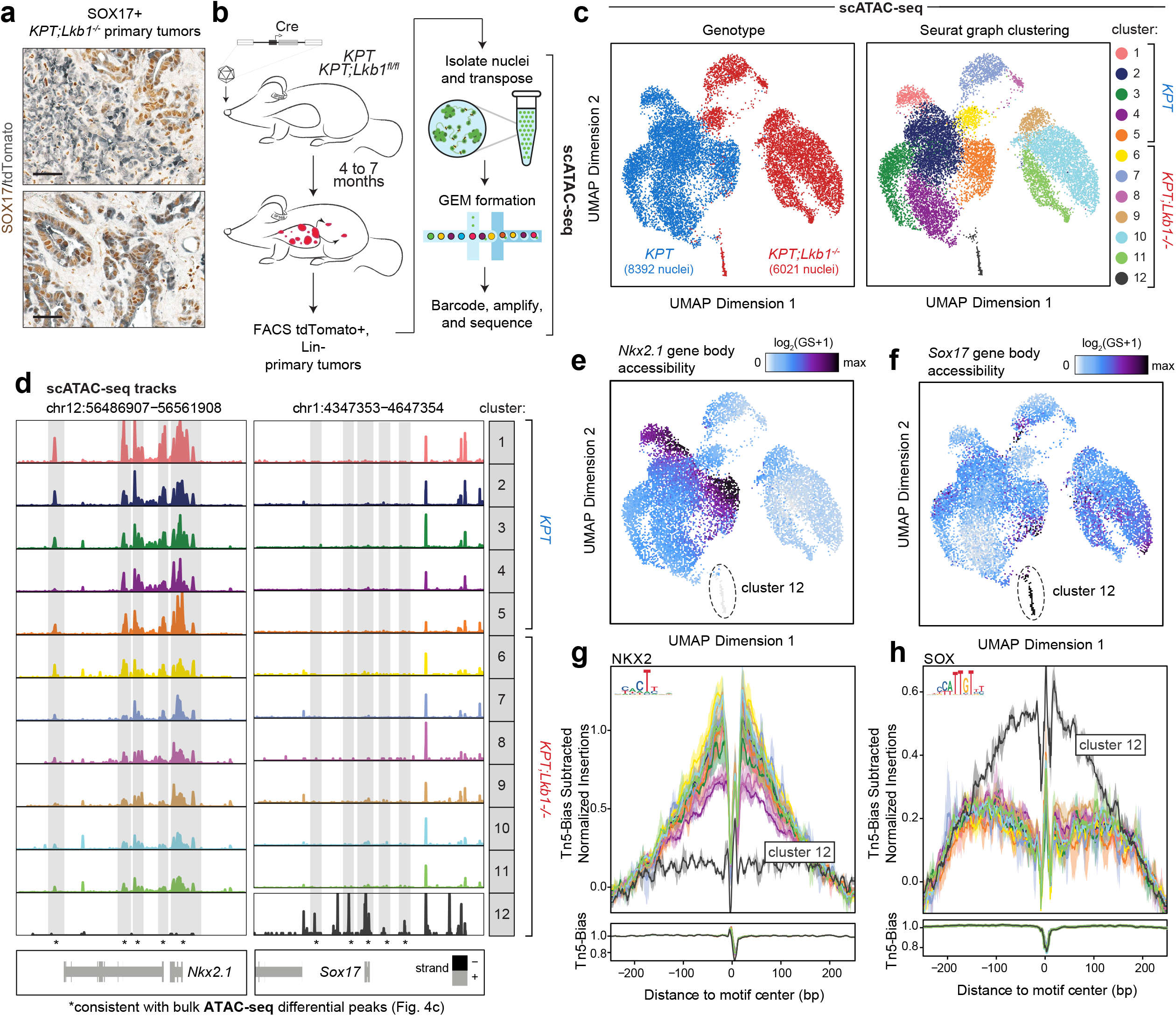
LKB1-deficient primary tumors harbor sub-populations of SOX17+ cells. **a.** Representative immunohistochemistry for SOX17 (in brown) and tdTomato (in grey) on two LKB1-deficient lung adenocarcinoma primary tumors. Scale bars represent 50uM. **b.** Schematic of tumor initiation and processing for scATAC-seq. tdTomato+, DAPI-cancer cells that were negative for the lineage (Lin) markers CD45, CD31, F4/F80, and Ter119 were sorted by FACS before scATAC-seq library preparation. **c.** Uniform Manifold Approximation and Projection (UMAP) of 8392 nuclei from 4 *KPT* primary tumors and 6021 nuclei from 3*KPT;Lkb1^−/−^* primary tumors, colored by genotype (*left*) or cluster according to Seurat graph clustering (*right*). **d.** *Nkx2.1* and *Sox17* genome accessibility tracks for each cluster indicated in Fig 4c. Significant ATAC-seq peaks from bulk chromatin accessibility profiling (Fig. 3e) are highlighted in grey and indicated with an asterisk (*).**e** and **f.** UMAP colored by the average gene body accessibility for *Nkx2.1* (**e**) or *Sox17* (**f**) in each cell. **g** and **h.** *Top*: Footprint of accessibility for each scATAC-seq cluster for genomic regions containing NKX2 (**g**) and SOX (**h**) motifs.*Bottom*: Modeled hexamer insertion bias of Tn5 around sites containing each motif.

To evaluate the epigenetic profiles of SOX17+ primary tumor cells, we performed droplet-based single-cell ATAC-seq (scATAC-seq)^31,32^ on cancer cells from LKB1-proficient (n=4) and LKB1-deficient (n=3) primary tumors (**Fig. 4b and Extended Data Figs. 9a, b**). We identified 12 distinct clusters of cells (**Fig. 4c** and **Extended Data Fig. 9c**; see Methods)^33^, with clusters 1-5 primarily composed of LKB1-proficient cells and clusters 6-12 primarily composed of LKB1-deficient cells. However, cells in cluster 12 (n=112 cells) stood out as a potential source of metastatically competent LKB1-deficient cells, exhibiting the highest accessibility near the *Sox17* locus as well as the lowest accessibility near the *Nkx2.1* locus (**Figs. 4d-f**). Cluster 12 is primarily composed of cells from two LKB1-deficient primary tumors derived from mouse 13 (13A and 13B). Motif enrichment and transcription factor footprinting^25^ revealed high flanking accessibility of SOX-containing genomic regions and a loss of the NKX2 footprint in cells in cluster 12 (**Fig.4g, h** and **Extended Data Fig. 9d**). Furthermore, genomic regions with the highest accessibility in LKB1-deficient primary tumors had the lowest average accessibility in cells in cluster 12 compared to clusters 1-11 (**Extended Data Fig. 9e**), while genomic regions with the highest accessibility in LKB1-deficient metastases had the highest average accessibility in cells in cluster 12 compared to clusters 1-11 (**Extended Data Fig. 9f**). Thus, sub-populations of cancer cells within LKB1-deficient primary tumors exhibit chromatin features suggestive of a SOX17+, metastatic-like state.

### SOX17 maintains accessibility of genomic regions containing SOX binding sites in metastatic, LKB1-deficient cells

To further establish a link between LKB1 and SOX17 during metastatic progression, we evaluated the effect of LKB1 restoration on SOX17 expression in our metastatic, LKB1-restorable cell lines (LR1 and LR2). Restoring LKB1 was sufficient to dramatically reduce *Sox17* mRNA expression and local accessibility at cis-regulatory sites near the *Sox17* locus (**Fig. 5a and Extended Data Fig. 10a, b**). Restoring LKB1 was also associated with a global loss of accessibility at genomic regions containing SOX binding sites in human and murine cell lines following LKB1 restoration (**Fig. 1d and Extended Data Fig. 2h-i, 6f**). GO term enrichment analysis of the genes closest to these genomic regions revealed decreased accessibility near genes related to the positive regulation of epithelial cell adhesion and extracellular matrix assembly, with implications for how cancer cells interact with the microenvironment and surrounding cell types (**Extended Data Fig. 10c**). Notably, inactivating the SIK family of kinases prior to LKB1 restoration was sufficient to maintain high *Sox17* mRNA expression and local chromatin accessibility at the *Sox17* locus (**Extended Data Fig. 10d, e**). These results suggest that LKB1-deficient metastases not only express higher levels of SOX17 compared to LKB1-proficient metastases, but also that the LKB1-SIK pathway actively inhibits the expression and thus activity of SOX17.

**Figure 5.**
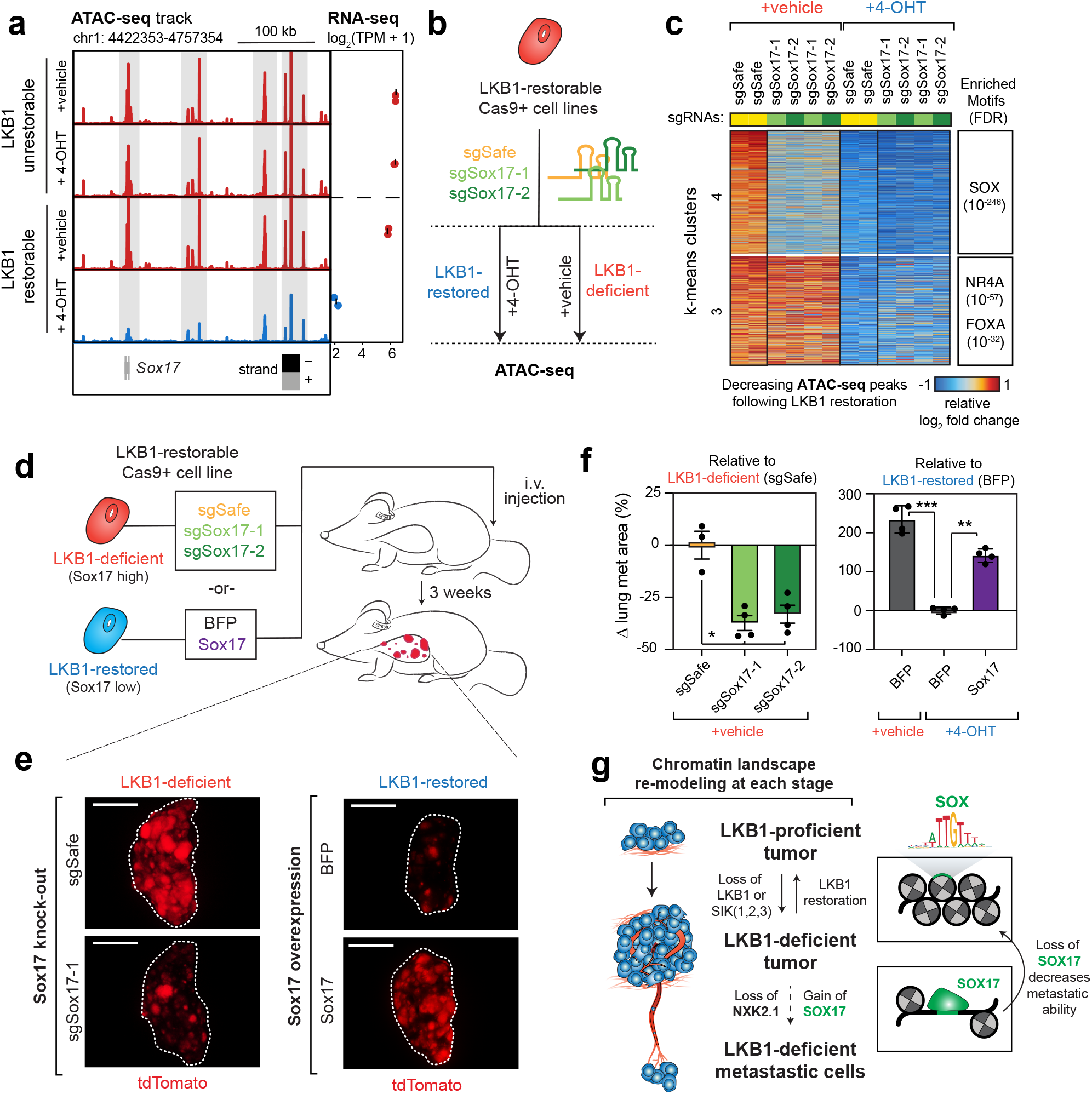
SOX17 regulates chromatin state and growth of metastatic, LKB1-deficient cells. **a.** *Sox17* genome accessibility track (*left*) and mean *Sox17* mNA expression across technical replicates (*right*) of an LKB1-unrestorable cell line (LU2) and a metastatic LKB1-restorable cell line (LR2) treated with 4-OHT or vehicle for sixdays. Highlighted in grey are significantly differential ATAC-seq peaks (log_2_ fold change < −0.5, FDR < 0.05) following LKB1 restoration. *Sox17* also has significantly decreased mRNA expression (log_2_ fold change < −1, FDR < 0.05) following LKB1 restoration. **b.** Schematic of knocking out the transcription factor Sox17 with and without LKB1 restoration in LKB1-restorable cell lines (LR1;Cas9 and LR2;Cas9) and performing ATAC-seq. **c.** Heatmap of the relative log_2_ fold changes in k-means clusters 3 and 4 of the indicated genotypes of cells with and without LKB1 restoration compared to the average log_2_ fold changes in sgSafe control cells. A subset (5,379 peaks; all decreasing peaks) of the top 10,000 consistent, variable ATAC-seq peaks following LKB1 restoration in cells transduced with either sgSafe or blue fluorescent protein (BFP) controls are shown. Full heatmap is shown in Extended Data Fig. 7f. **d.** Schematic of injecting LKB1-deficient cells expressing sgRNAs targeting Sox17 (sgSox17-1 and sgSox17-2) or injecting LKB1-restored cells expressing Sox17 cDNA intravenously (i.v.) into immunocompromised NSG mice. Tumor burden was analyzed three weeks post-injection. **e.** Representative fluorescent tdTomato+ images of single lung lobes following i.v. injection. **f.** Change in % tumor area compared to sgSafe + vehicle (*left*) or compared to BFP + 4-OHT (*right*). Each point represents an individual mouse, mean +/− SEM is shown. *p < 0.01, ** p < 0.001, *** p < 0.0001. **g.** Summary of LKB1-induced chromatin accessibility changes in lung adenocarcinoma primary tumors and metastases.

To evaluate whether SOX17 is required to maintain accessibility at genomic regions containing SOX binding sites, we inactivated *Sox17* with two sgRNAs in the LR2;Cas9 cell line and performed ATAC-seq with and without LKB1 restoration (**Fig. 5b and Extended Data Fig. 10f**). In LKB1-deficient metastatic cells, *Sox17* inactivation decreased accessibility at SOX-containing genomic regions to levels approaching that of LKB1-restored cells (**Fig. 5c and Extended Data Fig. 10g**). Next, we overexpressed *Sox17* cDNA and performed ATAC-seq with and without LKB1 restoration (**Extended Data Fig. 10h**). In LKB1-restored cells, *Sox17* expression led to the maintenance of accessibility at genomic regions containing SOX binding sites (**Extended Data Figs. 10i**; cluster 4). We confirmed these results in a second independent cell line (LR1;Cas9) (**Extended Data Fig. 10i**). Furthermore, expression of a sgRNA-resistant *Sox17* cDNA abrogated the effects of knocking out endogenous *Sox17* (**Extended Data Fig. 10i**). Therefore, SOX17 is necessary and sufficient to maintain accessibility at genomic regions containing SOX binding sites in LKB1-deficient, metastatic cells.

### SOX17 drives tumor growth of metastatic, LKB1-deficient cells

To further evaluate whether SOX17 regulates the growth of metastatic lung cancer cells, we inactivated *Sox17* with sgRNAs or overexpressed *Sox17* cDNA in the LR2;Cas9 LKB1-restorable cell line, restored LKB1, and injected each cell population intravenously into recipient mice (**Fig. 5d**). After three weeks of growth, we evaluated the colonization and growth of cells in the lung. Knocking out *Sox17* in LKB1-deficient cells resulted in a significantly reduced tumor burden relative to an sgSafe control (**Fig. 5e, f and Extended Data Fig. 11a**). In contrast, overexpressing *Sox17* in LKB1-restored cells increased tumor burden (**Fig. 5e, f and Extended Data Fig. 11b**). In addition, to evaluate the ability of *Sox17*-overexpressing cells to both leave the primary tumor and establish metastases, we injected LKB1-restored cells with and without overexpressed *Sox17* cDNA subcutaneously into recipient mice (**Extended Data Fig. 11c**). After five weeks of growth, we evaluated cells that had left the subcutaneous “primary” tumor and colonized in the lung(**Extended Data Fig. 11d**). While overexpressing *Sox17* did not change subcutaneous tumor growth (**Extended Data Fig. 11e**), *Sox17*-overexpressing cells had a significantly greater ability to colonize the lung (**Extended Data Fig. 11f**). Further, to evaluate the ability of *Sox17*-overexpressing cells to form metastases elsewhere in the body, we injected LKB1-restored cells with and without overexpressed *Sox17* cDNA intrasplenically into recipient mice (**Extended Data Fig. 11g**). After three weeks of growth, *Sox17-*overexpressing cells had a greater ability to colonize to the liver (p=0.055) (**Extended Data Fig. 11h-i**). Thus, SOX17 drives a genotype-specific epigenetic program that promotes the metastatic competency of LKB1-deficient cells.

## Discussion

Here we show that inactivation of LKB1, a tumor suppressive kinase, drives widespread chromatin accessibility changes in lung adenocarcinoma that evolve throughout cancer progression. While LKB1 has been well-studied for its metabolic roles in cancer, LKB1-induced chromatin changes are surprisingly AMPK-independent and depend almost exclusively on expression of the SIK family of kinases. Recent studies have additionally revealed that deleting AMPK hurts rather than helps lung cancer growth, and AMPK1 is preferentially amplified in lung adenocarcinoma, suggesting that AMPK is not a classic tumor suppressor in this cancer type^5,33^. Therefore, SIKs are emerging as the main drivers of LKB1-mediated tumor suppression and epigenetic regulation in lung cancer. Interestingly, the SIK family of kinases has a known role in the inhibition of class IIa histone deacetylases (HDACs)^34^. In contrast to other classes of HDACs, Class IIa HDACs do not have the typical core enzymatic domain required for deacetylating histones; however, they form multiprotein complexes with transcription factors to interact with chromatin^35^. Thus, the SIK-HDAC relationship might be relevant for future studies attempting to dissect the regulation of SOX17 and the overall chromatin accessibility states of LKB1-deficient and LKB1-proficient cells.

Overall, our findings reveal that inactivation of LKB1/SIK signaling drives two separate waves of epigenetic re-modeling, with the first set of changes occurring within lung primary tumors and the second set of changes mediated by cis-regulatory activation of the transcription factor SOX17 in metastatic cells (**Fig. 5g**). Thus, the downstream effects of a driver mutation can change throughout tumor development and subsequently enhance metastatic ability. While the LKB1 pathway likely constitutively represses SOX17, the consequences of this repression are not observed until SOX17 expression is activated during metastatic transformation. However, as not all LKB1-deficient cancer cells express SOX17, there must be a second signal that initiates the metastatic program, which currently remains unknown. Regulation of the strong endodermal transcription factor SOX17 could also have implications for the diverse histological sub-types observed in LKB1-deficient lung tumors^8,36^, and further work to understand the plasticity of LKB1-deficient cells in the context of such widespread chromatin accessibility changes is warranted.

By resolving the epigenetic landscape of lung adenocarcinoma primary tumors at single-cell resolution, we further discovered sub-populations of cancer cells in primary tumors that share a common epigenetic state with the cancer cells in metastases. This result suggests that primary tumors harbor rare and epigenetically distinct cells that are “poised” to seed distant metastases, rather than evolving a specialized cell state after metastatic colonization. An early mechanism of epigenetic transformation opens up the possibility of identifying biomarkers to predict which tumors have already seeded micrometastases before detection is possible. In addition, we anticipate that genotype-driven epigenetic differences between primary tumors and metastases will likely inform how patients respond to personalized therapies. Overall, these findings help to explain the paradox wherein primary tumors and metastases share the same genetic mutations yet exhibit extremely different behaviors, and we anticipate that an evolving mechanism of tumor suppression is more broadly applicable to other commonly mutated driver genes and cancer types.

## Materials and Methods

### Murine cell lines

Murine cell lines were generated from individual primary tumors and metastases from *Kras^LSL-G12D^;Trp53^flox/flox^;Lkb1^XTR/XTR^;Rosa26^LSL-tdTomato^* (cell lines LU1 and LU2), *Kras^LSL-G12D^;Trp53^flox/flox^;Lkb1^XTR/XTR^;Rosa26^FlpOER/LSL-tdTomato^* (cell line LR2), and *Kras^LSL-G12D^;Trp53^flox/flox^;Lkb1^XTR/XTR^;Rosa26^FlpOER/+^* (cell line LR1) mice previously transduced with lentiviral Cre. The *Lkb1^XTR/XTR^* mouse allele is currently unpublished but was generated using the same design and methods as outlined for the *Tp53^XTR/XTR^* allele^19^. Sequences of the allele will be made available on request. All cell lines have gene expression patterns consistent with being in a metastatic state (*Nkx2.1^low^*;*Hmga2^high^*) (**Supplementary Table 3**). To derive cell lines, tumors were excised from the lungs or lymph nodes of mice, minced into pieces using scissors, and directly cultured in DMEM media supplemented with 10% FBS, 1% penicillin-streptomycin-glutamate, and 0.1% amphotericin at 37°C with 5% CO_2_ until cell line establishment. Cells were authenticated for genotype. All human cell lines tested negative for mycoplasma using the MycoAlert Mycoplasma Detection Kit (Lonza).

All four murine cell lines (LR1, LR2, LU1, and LU2) were grown in DMEM media supplemented with 10% FBS, 1% penicillin-streptomycin-glutamate, and 0.1% amphotericin. LR1 and LR2 cell lines were then transduced with an SpCas9 lentiviral vector with a Blasticidin selection marker (Addgene #52962) and selected with Blasticidin (10ug/mL) for >5 days. To be able to test Cas9 cutting efficiency, site-directed mutagenesis was used to delete a loxP site in the pMCB306 backbone (Addgene #89360), since these cell lines were previously transduced with Cre recombinase to initiate tumor growth in mice. This plasmid is a self-GFP cutting reporter with both expression of GFP and a sgRNA against GFP on the same backbone. Polyclonal Cas9+ populations with high cutting efficiency were established and used for subsequent experiments (referred to as LR1;Cas9 and LR2;Cas9 in the text). For LKB1 restoration induction, cells were treated with either 1uM 4-hydroxytamoxifen (4-OHT; Sigma Aldrich) dissolved in 100% ethanol or a vehicle (1:2000 100% ethanol) for the indicated time-points.

### Proliferation doubling assays

For population doubling assays, cell lines were treated with 4-OHT or vehicle for twelve days. Every other day, cells were trypsinized for 5 minutes at 37°C, collected in microcentrifuge tubes, counted, and re-plated with 50,000 cells per well of a 6-well in triplicate. The total number of cells in each well was recorded for each day. The number of population doublings was assessed by taking the total number of cells (N) for that day and normalizing to the original 50,000 cells plated i.e. log_2_(N/50000). Cumulative population doublings are the sum of the population doublings from the current time-point as well as all previous time-points. Two-tailed t-tests were performed to determine statistical significance.

### Clonogenic growth assays

For clonogenic growth assays, cell lines were pre-treated with a vehicle control or 4-OHT for six days. Cells were trypsinized for 5 minutes at 37°C, collected in conical tubes, counted, and re-plated at 500 cells/well of a 6-well plate in triplicate. Plates were incubated at 37°C with 5% CO_2_ for six days. For analysis, cells were rinsed with room temperature PBS, fixed with ethanol for 5 minutes at room temperature, and stained with 1% crystal violet solution in water (Millipore-Sigma) for an additional 5 minutes. Plates were rinsed with water and left to dry overnight, followed by scanning into the computer and analysis using ImageJ. The % area of the plate covered by cells was normalized to the average % area of the plate covered by cells treated with a vehicle control. Two-tailed t-tests were performed to determine statistical significance.

### RNA-sequencing library preparation for cell lines

Cell lines were treated with 4-OHT or vehicle for six days prior to RNA extraction. Adherent cells were rinsed with PBS, trypsinized for 5 minutes at 37°C, spun down, and cell pellets were frozen at −80°C. Cell pellets were processed to total RNA using the RNeasy Plus Mini Kit (Qiagen) according to standard protocols. RNA quality was assessed using the Bioanalyzer 2100 Agilent). All of the RNA used for RNA-seq had an RNA integrity number (RIN) of 10.0. 500ng total RNA for each sample was processed into libraries using the TruSeq RNA Library Prep Kit v2 (Illumina) and sequenced according to standard protocols.

### RNA-sequencing data processing and alignment

RNA-seq data was first trimmed with CutAdapt and then aligned with kallisto^37^. We downloaded pre-compiled transcriptome indices from https://github.com/pachterlab/kallisto-transcriptome-indices/releases for mm10 and hg38. We aligned with kallisto quant using the following parameters: “kallisto quant –genomebam –gtf –chromosomes –threads –index”. This generated a transcript count file that was converted to gene counts using tximport. We then created a SummarizedExperiment in R containing a matrix of the samples by genes with the gene coordinates. We used the genomebam created by kallisto to validate the number of reads per exon in LKB1 (the trapped configuration of the XTR allele causes early termination of transcription after exon 1, **Extended Data Fig. 1d**).

### RNA-seq data analysis – Differential Expression

To compute differential gene expression, we used edgeR’s glmQLFTest. We used as input two groups with a simple design with a 0 intercept “~0 + Group”. We first calculated normFactors using the TMM normalization “calcNormFactors(y, method = “TMM”)”. Next, we estimated dispersions with robustness “estimateDisp(y, design = design, robust = TRUE). Then we fitted the generalized linear model using “glmQLFit(y, design = design)”. Lastly, we used the glmQLFTest to compute log_2_ fold changes and adjusted p-values. We chose the indicated significance cutoffs based on the thresholds set by our control LKB1-unrestorable cell lines (LU1 and LU2) treated with 4-OHT.

### Immunoblot analysis

Adherent cells were rinsed with ice-cold PBS, lysed in RIPA buffer, scraped from plates, and spun at 13,000g for 30 minutes at 4°C. The concentration of protein-containing supernatant was quantified using the Pierce BCA Protein Assay Kit (Thermo Fisher). 10ug of each sample was loaded onto NuPage 4-12% Bis-Tris protein gels (Thermo Fisher) and transferred to polyvinylidene fluoride (PVDF) membranes (Bio-Rad) at 10V overnight. Blocking, primary, and secondary incubations were performed in Tris-buffered saline (TBS) with 0.1% Tween-20. Blocking was performed in 5% dry milk and primary antibody incubation was performed in 5% bovine serum albumin (BSA) (Cell Signaling). Secondary antibody incubation was performed in 5% dry milk with anti-rabbit (Cell Signaling, 7074S) or anti-mouse (BD Biosciences, 610418) antibodies. LKB1 (Cell signaling, 13031S) and SOX17 (Abcam ab224637) protein expression was assessed by Western blotting. HSP90 (BD Biosciences, 610418) was used as a sample processing control on a separate blot that was processed in parallel with the same input master mix. Full blots are shown in **Extended Data Figure 12**.

### Allograft studies in immunocompromised mice

For intravenous transplants into immunocompromised NSG mice, cells were treated with either a vehicle control or 4-OHT for six days and 5 × 10^4^ cells were injected into one of lateral tail veins. Mice were sacrificed 21 days post-injection. For intrasplenic transplants into immunocompromised NSG mice, cells were treated with 4-OHT for six days and 5 × 10^4^ cells were injected via intrasplenic injection. To perform intrasplenic injections, the left flank of each mouse was shaved and disinfected with 70% ethanol. A small incision was made to expose the spleen and a ligation on the splenic branch of the lienopancreatic artery was performed. Following injection of cells, a surgical knot was made in the upper part of the spleen and the lower part of the spleen was removed prior to sewing the body wall back with surgical knots. The skin incision was closed with staples and antiseptic solution was applied to clean the wound. Mice were sacrificed three weeks post-injection. For subcutaneous transplants into immunocompromised NSG mice (**Extended Data Fig. 1**), 2 × 10^5^ untreated cells were re-suspended in 200uL PBS and injected into two sites per mouse. Once tumors were readily palpable, mice were randomized and treated via oral gavage with either a vehicle control (200uL 10% ethanol 90% corn oil) or tamoxifen (200uL of 20 mg ml^−1^ tamoxifen dissolved in 10% ethanol 90% corn oil) (Sigma Aldrich) for three consecutive days. Tamoxifen is metabolized to 4-OHT in the liver and systemically distributed throughout the body. The height, width, and length, of each tumor was measured using calipers every two days for 14 days (LU1, LR2) or every four days for 16 days (LU2, LR1). Tumor volume was roughly calculated by multiplying height × width × length of each tumor. During the experimental time-course, subcutaneous tumors never grew above 1.0 cm^3^. For subcutaneous transplants to model metastatic spread to the lung, 5 × 10^4^ cells of the indicated genotypes pre-treated with six days of 4-OHT or vehicle were re-suspended in 200uL PBS and injected into two sites per mouse. Mice were sacrificed five weeks post-injection. The Stanford Institute of Medicine Animal Care and Use Committee approved all animal studies and procedures.

### Immunohistochemistry and histological quantification

Lung samples were fixed in 4% formalin and paraffin embedded. Hematoxylin and eosin staining was performed using standard methods and percent tumor area was calculated using ImageJ. For IHC, we used an antibody to SOX17 (Abcam, ab224637) at a 1:1000 dilution. Heat-mediated antigen retrieval was performed in Tris/EDTA buffer with pH 9.0. To evaluate SOX17 expression, we quantified the number of tumors with tumor area composed of 0% SOX17+ cells (none), <25% SOX17+ cells (low), 25-50% SOX17+ cells (medium), and >50% SOX17+ cells (high) using ImageJ.

### Lentiviral production

All lentiviruses were produced by co-transfecting lentiviral backbones with packaging vectors (delta8.2 and VSV-G) into 293T cells using PEI (Polysciences). The viral-containing supernatant was collected at 48- and 72-hours post-transfection, filtered through a 0.45uM filter, and combined with fresh media to transduce cells. Cells were exposed to viral-containing supernatant for up to two days prior to the first fresh media change. Human cell lines were also incubated with 8ug/mL polybrene (Sigma) to enhance transduction efficiency.

### CRISPR/ Cas9 screen and sample processing

The genome-scale CRISPR/Cas9 knock-out library was synthesized by Agilent and designed and cloned as previously described^20^. The genome-scale library was designed to have ~200,000 sgRNAs targeting ~20,000 coding genes (10 sgRNAs per gene), with >13,000 negative control sgRNAs that are either non-targeting (sgNT) or safe-targeting (sgSafe) (**Supplementary Table 1**). This library is composed of ten sub-library pools roughly divided according to gene function (https://www.addgene.org/pooled-library/bassik-mouse-crispr-knockout/). The entire genome-scale screen was performed in two halves, each composed of five sub-library pools. In addition, the second half of the screen included a repeat of the sub-library containing sgRNAs targeting *Lkb1* as a positive control. The two screens were performed sequentially.

For both halves of the screen, the combined sub-library plasmid pools were transfected into 293T cells to produce lentiviral pools, which were transduced into LR1;Cas9 cells. Cells were transduced at a multiplicity of infection of 0.3, and after 48 hours were selected with puromycin (8 ug/mL) for 3 days until the library-transduced population was >90% mCherry+ (a marker for lentivirus transduction). Cells were expanded for another 2 days and aliquots were saved as day 0 stocks in liquid nitrogen. Remaining cells were plated and treated in duplicate with either vehicle or 4-OHT. To maintain library complexity, the screens were performed at 200x cell number coverage per sgRNA. Due to the fast doubling time of this cell line, each half of the screen required passaging >165 15cm dishes every two days. 12 days later, cells were collected and stored in cryovials in liquid nitrogen for further processing. Genomic DNA was extracted from each sample in technical duplicate with the Qiagen Blood Maxi Kit (Qiagen). sgRNA cassettes were PCR-amplified from genomic DNA and sample indices, sequencing adapters, and flow-cell adapters were added in two sequential rounds of PCR as previously described^20^.

### CRISPR/Cas9 screen data alignment and analysis

We aligned each half of the genome-scale CRISPR/Cas9 screen individually using casTLE^38^, which uses bowtie alignment. This alignment returned a counts matrix for each sgRNA per sample. We then identified the sgRNAs that were overlapping in each half of the CRISPR screen and then computed the mean reads in these sgRNAs. We then scaled each half of the screen such that the mean reads in overlapping sgRNA was identical. We then used the values from each half for sgRNAs specific to that half and overlapping sgRNAs the mean reads across both screen halves was used. We then depth-normalized across all samples. We computed the log_2_ correlations and plotted the Pearson correlation matrix in R. To quantify the sgRNAs that were enriched/disenriched in the screen we used MAGeCK^39^. Briefly, we used mageck test with parameters “-k counts.tsv -t day12_LKB1_Restored -c day12_LKB1_Unrestored”. We then accessed the MAGeCK RRA scores from gene_summary.txt file and filtered targets with less than 5 sgRNAs assigned to each target. We then took the top 50 sgRNA and used them as input to PANTHER GO term enrichment.

### ATAC-sequencing library preparation for cell lines

Cell lines were treated with 4-OHT or vehicle for the indicated time-points prior to transposition. For the ATAC-seq time-course (Extended Data Fig. 2), samples were treated in a reverse time-course such that transposition for all time-points occurred at the same time. The media for all cells was changed at each time-point to control for fluctuations in growth factors or other media contents between samples. For all experiments, adherent cells were rinsed with PBS, trypsinized for 5 minutes at 37°C, spun down, and re-suspended in PBS. After the cells were counted, 50,000 cells in technical duplicate were resuspended in 250uL PBS and centrifuged at 500rcf for 5 minutes at 4°C in a fixed-angle centrifuge. Pelleted cells were re-suspended in 50uL ATAC-seq resuspension buffer (RSB; 10mM Tris-HCl pH 7.4, 10mM NaCl, and 3mM MgCl_2_ in ddH_2_O made fresh) containing 0.1% NP40, 0.1% Tween-20, and 0.01% digitonin according to the omni-ATAC-seq protocol^23^. After incubating on ice for 3 minutes, 1mL of ATAC-seq RSB containing 0.1% Tween-20 was added. Nuclei were centrifuged at 500rcf for 5 minutes at 4°C in a fixed-angle centrifuge, 900uL of the supernatant was taken off, and then the nuclei were centrifuged for an additional 5 minutes under the same conditions. The remaining 200uL of supernatant was aspirated and nuclei were resuspended in 50uL of transposition mix (25uL 2X TD buffer (2mL 1M Tris-Hcl pH 7.6, 1mL 1M MgCl_2_, 20mL DMF, and 77mL ddH_2_O aliquoted and stored at −20°C), 2.5uL transposase (100nM final), 16.5uL PBS, 0.5uL 1% digitonin, 0.5uL 10% Tween-20, and 5uL ddH_2_O). Transposition reactions were incubated at 37°C for 30 minutes with 1,000 r.p.m. shaking in a thermomixer and cleaned up using MinElute PCR purification columns (Qiagen). The transposed samples were then amplified to add sample indices and sequencing flow cell adapters and cleaned up as described previously, with a target concentration of 20uL at 4nM. Paired-end sequencing was performed on an Illumina NextSeq using 75-cycle kits.

### ATAC-seq data processing and alignment

Adaptor sequence trimming, mapping to the mouse (mm10) or human (hg38) reference genome using Bowtie2 and PCR duplicate removal using Picard Tools were performed. Aligned reads (BAM) mapping to “chrM” were also removed from downstream analysis. BAM files were subsequently corrected for the Tn5 offset (“+” stranded +4 bp, “−” stranded −5 bp) using Rsamtools “scanbam” and Genomic Ranges. These ATAC-seq fragments were then saved as R binarized object files (.rds) for further downstream analysis.

### ATAC-seq data QC – Transcription start site enrichment

Enrichment of ATAC-seq accessibility at transcription start sites (TSSs) was used to robustly quantify ATAC-seq data quality without the need for a defined peak set as previously described^25^. First, ATAC-seq fragments were read into R with “readRDS”. To get the TSS enrichment profile, each TSS from the R package “TxDb.Mmusculus.UCSC.Mm10.knownGene” or “TxDb.Hsapiens.UCSC.hg38.knownGene” (accessed by transcripts(TxDb)) were extended 2000 bp in each direction and overlapped with the insertions (ends of each ATAC-seq fragment from above) using “findOverlaps”. Next, the distance between the insertions and the strand-corrected TSS was calculated and the number of insertions occurring in each single-base bin was summed. To normalize this value, we divided the accessibility at each position +/− 2000 bp from the TSS to the mean of the accessibility at flanking positions +/−1900-2000 bp from the TSS. The final TSS enrichment reported was the maximum enrichment value within +/− 50 bp of the TSS after smoothing with a rolling mean every 51 bp.

### ATAC-seq data analysis – Creating a High-Quality ATAC-seq Peak Set for Experiments in Murine Cell Lines

Peak calling for all ATAC-seq profiles was performed to ensure high quality fixed-width peaks as described previously ^25^. For each sample, peak calling was performed on the Tn5-corrected single-base insertions (ends of each ATAC-seq fragment from above) using the MACS2 callpeak command with parameters “–shift −75 –extsize 150 –nomodel –call-summits –nolambda –keep-dup all -q 0.01”. The peak summits were then extended by 250 bp on either side to a final width of 501 bp, filtered by the ENCODE hg38/mm10 blacklist (https://www.encodeproject.org/annotations/ENCSR636HFF/), and filtered to remove peaks that extend beyond the ends of chromosomes.

Overlapping peaks called within a single sample were handled using an iterative removal procedure as previously described^25^. We also modified the MACS2 normalization to a quantile normalization since we used qvalues (“−q” param) in MACS2 vs the depth normalization. We then tested the reproducibility for each peak by overlapping peak calls from individual replicates with this peak set to identify which were called more than one time and kept these peaks. We then combined all peak calls from all samples within an experiment and performed the iterative removal procedure again. Lastly, any peaks that spanned a genomic region containing “N” nucleotides and any peaks mapping to the Y chromosome were removed. This resulted in a set of high quality, reproducible, fixed-width peaks for each experiment.

Throughout this project we continually performed experiments and continuously re-establishing a common peak set across all experiments became unreasonable. Creating a peak set across all experiments also leads to the addition of peaks that are not relevant to the current experiment which can be problematic when doing many orthogonal perturbations across disparate experiments. Therefore, we developed a new approach for using a “basis experimental peak set” to facilitate easier cross comparisons. This meant that after getting a non-overlapping fixed-width peak set per experiment we identified which peaks overlapped our “basis experimental peak set” and replaced those peaks with the exact coordinates of the “basis experimental peak set”. The peaks being all uniform in size prevented large changes in these peak regions, but now provided peaks that were present across each experiment. Additionally, we tested the results with and without this procedure and the differences were minimal. This approach can help alleviate some of the challenges with having to re-define peak sets as you continually perform experiments based on previous experiments which then subsequently get affected with a new peak set. We chose the LKB1-restoration time course as our basis experimental peak set (**Extended Data Fig. 2**) it captures majority of the changes throughout LKB1 progression (LR1 and LR2) and was high quality.

### ATAC-seq data analysis – Differential Accessibility

To compute differential accessibility, we used edgeR’s glmQLFTest similarly to our RNA-seq. We used as input two groups with a simple design with a 0 intercept “~0 + Group”. We first calculated normFactors using the TMM normalization “calcNormFactors(y, method = “TMM”)”. Next, we estimated dispersions with robustness “estimateDisp(y, design = design, robust = TRUE). Then we fitted the generalized linear model using “glmQLFit(y, design = design)”. Laslty, we used the glmQLFTest to compute log_2_ fold changes and adjusted p-values. We chose the significance cutoffs based on our non-restorable control LKB1-experiment.

### ATAC-seq data analysis – chromVAR for transcription factor activity

We wanted to measure global TF activity using chromVAR. We used as input the raw counts for all peaks and the CIS-BP motif matches (from chromVARMotifs “mouse_motifs_v1”, see https://github.com/GreenleafLab/chromVARmotifs) within these peaks from motifmatchr. We then computed the GC bias-corrected deviations and deviation scores using the chromVAR “deviations” function.

We wanted to compare our results across species and experiments in a way that captured the downstream effectors in the LKB1-chromatin pathway. To do this comparison, we developed a way to test differences in TF chromVAR deviation scores across multiple experiments. Since there are multiple assigned motifs for many TFs, comparisons intra-species we compared the exact same motifs and inter-species we compared all combinations of the motifs. In our motif database, we found excess number of FOX related motifs (human >200 motifs for the FOX family and in mouse > 100 motifs), thus we limited all FOX motifs to those in FOXA1 and FOXA2 because our initial LKB1-restoration experiment showed changed in expression in those two FOX motifs. However, all q-values accounted for each total FOX combination to make sure that this decision didn’t influence our results. Once we had computed all the TF-TF comparisons across each experiment, we then computed the differences the two groups in each experiment. We then permuted the samples (n = 10,000) within each of the two groups in each experiment and computed the Euclidean distance from the origin (0,0). We then computed the number of times the permuted distance was greater than the distance in the original comparison to determine the p-value. This p-value was determined for all TF-TF combinations. We then converted the p-values to q-values using “qvalue” in R. We determined significantly differential TFs as those whose q-value < 0.05 and distance was greater than 5. Additionally, we required that the absolute difference in either comparison was greater than 1 to prevent differences that were only in 1 experiment. We mainly focused on TFs that were commonly different in the top-right and bottom-left quadrant. Additionally, we tried to highlight the same TF families across all the comparisons to show the consistency of these results. This comparison allows better interpretability of comparing relative differences vs peaks because chromVAR converts the accessibility data to TFs. Additionally, this comparison allows for quantitative TF difference comparisons while traditional differential testing with motif enrichment relies on comparing relative p-values in significantly differential peaks. Lastly, this comparison is normalized for GC-bias while differential testing with motif enrichment does not.

### ATAC-seq data analysis – Constructing a counts matrix and normalization

To get the number of ATAC-seq Tn5 insertions per peak, each Tn5 insertion site (each end of a fragment) was counted using “countOverlaps”. This was done by resizing each fragment to the start (“resize(fragments, 1, “start”)) and end (“resize(fragments,1,”end”)). This was done for all individual replicates and a counts matrix was compiled. From this, a SummarizedExperiment was constructed including peaks (rowRanges) as GenomicRanges, a counts matrix (assays), and metadata (colData) detailing information for each sample. The counts matrix was then normalized by using edgeR’s “cpm(matrix, log = TRUE, prior.count = 3)” followed by a quantile normalization using preprocessCore’s “normalize.quantiles” in R. The prior count is used to lower the contribution of variance from elements with lower count values. We used a prior count of 3 for ATAC-seq because there are more features with fewer relative counts than RNA-seq.

### ATAC-seq data analysis – Principal Component Analysis and K-means Heatmaps

To visualize our multi-dimensional data in reduced dimensions, we used principal component analysis (PCA) with “prcomp” in R. Prior to PCA we identified and then subsetted the top variable peaks (of the log2 normalized ATAC-seq matrix using matrixStats::rowVars), default as 10,000, to capture majority of the variance in the first few principal components. This procedure helps reduced batch effects and focuses on the biological variance in the data set. Additionally, we used these variable peaks as input for our heatmaps. Since there are a lot of features, it is efficient to use K-means clustering to group peaks with similar patterns for making a heatmap. By default, we chose to identify 5 k-means clusters and if we observed additional/less variance within these clusters we adjusted this number accordingly. We then used the peaks in each k-means to create a scaled log2 accessibility heatmap with ComplexHeatmap in R. This procedure enabled grouping peaks based on their accessibility patterns which could subsequently be used for motif hyper-geometric enrichment testing using the motifmatches from motifmatchr (see above).

### ATAC-seq data analysis – Comparisons with RNA-seq

To identify consistently differential genes (|LFC| > 0.5 and FDR < 0.01) that had consistent local accessibility changes (|LFC| > 0.5 and FDR < 0.05) we first identified differential peaks and genes (see above) for each restorable cell line. Next, we resized each gene coordinate from the start +/− 50 kb with resize(resize(gr, 1, “start”), 100001, “center”) in R. We then computed overlaps between these extended gene windows and the differential peaks. We then computed the average LFC across both cell lines for RNA and the number of consistent differential peaks within the extended gene window. We referred to these genes as “accessibility-linked genes”. These genes are higher candidate genes as they are consistently identified across different modalities.

To identify transcription factors that were enriched in the LKB1-restoration peaks, we computed the TPM’s for each gene and used the log_2_ fold changes. This analysis helps elucidate specific TFs within each family that are more likely to be the actual chromatin regulators. However, this analysis puts more emphasis on regulation through direct gene expression of the TF vs co-factors and other indirect measures. To identify which TFs within the NKX2 and SOX families were differentially regulated in the primary tumors and metastases we computed the gene body accessibility for each gene genome-wide. We then normalized the total accessibility to 1,000,000 counts and then log2 transformed the matrix. We then plotted at the differential gene body accessibility between the metastases and primary tumors for each of the NKX2 and SOX TFs. Additionally, we plotted the differential gene expression for these TFs and found strong agreement.

### ATAC-seq data Analysis – Reads-in-peaks-normalized bigwigs and sequencing tracks

To visualize our ATAC-seq data genome-wide we used ATAC-seq signal tracks that have been normalized by the number of reads in peaks as previously described ^25^. For reads-in-peaks normalization, we first constructed bigwigs based on the Tn5 insertion sites (ends of each fragment). To do this, the genome was binned into 100-bp intervals using “tile” in GenomicRanges of the chromosome sizes in R. The insertion sites (GenomicRanges) were then counted in each of these regions and then converted to a run-length encoding. We then normalized the total number of reads by a scale factor that converted all samples to a constant 10 million reads within peaks. This was then converted into a bigwig using rtracklayer “export.bw” in R. For plotting tracks, the bigwigs were read into R using rtracklayer “import.bw(as=”Rle”)” and plotted within R using ggplot2. All track figures in this paper show groups of tracks with matched y-axis scales.

### ATAC-seq data Analysis – TCGA LUAD Tumors

We downloaded the TCGA-ATAC-seq data from https://gdc.cancer.gov/about-data/publications/ATACseq-AWG for tumors with matched RNA. We then scored each tumor for being high (top 10%), medium (middle 80%) and low (bottom 10%) in LKB1 (STK11) expression (TPM). We additionally identified which tumors had a medium-high predicted mutation (VarScan2). We then unbiasedly identified the top 10,000 variable peaks and grouped them into 5 k-means clusters. We then plotted a heatmap of the scaled log2 accessibility as described above. To test the enrichment of specific gene mutations in each chromatin sub-type (See **Extended Data Fig. 4a**), we computed the proportion of medium-high predicted mutation burden of the gene (VarScan2) and computed a binomial enrichment vs the mutation frequency of all TCGA LUAD tumors (n = 230). We then computed the FDR for the binomial enrichments with p.adjust in R.

### scRNA-seq Analysis – LUAD Metastases Laughney et al. 2020

We downloaded the raw data from Laughney et al. 2020 from https://s3.amazonaws.com/dp-lab-data-public/lung-development-cancer-progression/PATIENT_LUNG_ADENOCARCINOMA_ANNOTATED.h5. We then read the subgroup (hdf5 formatted file) “INDF_EPITHELIAL_NOR_TUMOR_MET” for the normalized scRNA-seq matrix. We then computed z-scores for all genes. We then averaged the scaled expression for all cells from each donor that belonged in cluster “H0” and “H3” (to increase number of donors) which represent the most undifferentiated metastatic cells. We then computed the standard error of the mean (SEM) for all cells from each donor in these clusters. We then plotted the average and SEM SOX17 expression vs STK11 (LKB1) expression for each of the donors.

### Cloning and generating knock-out and overexpression cell lines

To generate individual knock-out cell lines, we first cloned individual sgRNAs into the pMJ114 backbone (Addgene #85995) using Q5 site-directed mutagenesis (NEB). A list of all sgRNA sequences used in this study is located in **Supplementary Table 4**. sgRNA sequences were chosen based on the most highly enriched sgRNAs in the genome-scale screen (sgLkb1) or by choosing the top two sgRNAs with the highest predicted cutting activity from the Brie library on Addgene (#73633). After making lentivirus and transducing cells with the lentiviral supernatant, we waited two days and then selected cells with 8ug/mL puromycin for at least three days to enrich for cells transduced with the lentivirus, before initiating treatment with vehicle or 4-OHT.

To generate double and triple knock-out cell lines, we used Gibson assembly to create lentiviral vectors with sgRNAs transcribed in series by the bovine U6 promoter, human U6 promoter, and murine U6 promoter, as previously described ^40^. In brief, we first cloned individual sgRNAs into the pMJ114 (Addgene #85995), pMJ117 (Addgene #85997), and pMJ179 (Addgene #85996) backbones, then stitched them together using Gibson assembly (NEB). For LKB1 downstream effector families with only two gene paralogs, we still included the third murine U6 promoter driving expression of sgSafe-1 to control for the effects of three cutting events occurring simultaneously in the same cell. Similarly, for the sgLkb1 control experiments, a bovine U6 promoter driving expression of sgLkb1-1 was combined with a human and mouse U6 driving expression of sgSafe-1 and sgSafe-2. After transducing cells with the lentiviral supernatant, we waited two days and then selected cells with 8ug/mL puromycin for at least three days to enrich for cells transduced with the lentivirus, before initiating treatment with vehicle or 4-OHT.

To generate cell lines with overexpression of Sox17, we codon optimized murine Sox17 cDNA to simultaneously mutate the sgSox17-1 and sgSox17-2 cut sites and ordered this sequence as a gBlock (IDT). We used Gibson assembly to replace BFP in pMJ114 with this modified Sox17 sequence. After making lentivirus and transducing cells with the lentiviral supernatant, we waited two days and then selected murine cell lines with 8ug/mL puromycin for at least three days to enrich for cells transduced with the lentivirus, before initiating treatment with vehicle or 4-OHT.

To generate human cell lines with overexpression of KEAP1 or LKB1, we amplified human KEAP1 and LKB1 off of human cDNA and used Gibson assembly to replace GFP in pMCB306 (Addgene #89360) with these sequences. After making lentivirus and transducing human cells with the lentiviral supernatant, we waited 2 days, selected human cell lines with 2ug/mL puromycin for at least 4 days to enrich for cells transduced with lentivirus, then let the cells recover in fresh media for 2 days before collecting for ATAC-seq library preparation.

### Human cell lines

All human non-small cell lung cancer cell lines (NCIH1437, A549, NCIH460, NCIH1355, NCIH1650, NCIH1975, NCIH358, NCIH2009) were either purchased directly from ATCC or were a gift from Dr. Michael Bassik’s laboratory, who previously purchased them from ATCC. Human cell lines were cultured in RPMI media supplemented with 10% FBS, 1% penicillin-streptomycin-glutamate, and 0.1% amphotericin. All human cell lines tested negative for mycoplasma using the MycoAlert Mycoplasma Detection Kit (Lonza).

### Autochthonous mouse models

Homozygous floxed Lkb1 alleles (*Lkb1^fl/fl^*) were bred into the metastatic KPT (Kras^LSL-G12D^;p53^fl/fl^;Rosa26^LSL-tdTomato^) model to generate LKB1-proficient and LKB1-deficient models of lung adenocarcinoma metastasis. Lentiviral Cre recombinase was co-transfected with packaging vectors (delta8.2 and VSV-G) into 293T cells using PEI, the supernatant was collected at 48 and 72 hours post-transfection, ultracentrifuged at 25,000rpm for 90 minutes, and resuspended in PBS. Tumors were initiated by intratracheal transduction of mice with lentiviral vectors expressing Cre recombinase, as previously described^41^. For ATAC-seq, tumors were collected and processed at staggered time-points where tumor burden was similar between *KPT* and *KPT;Lkb1^fl/fl^* cohorts of mice. For the survival curve, mice were sacrificed immediately after exhibiting physical symptoms of distress from lung tumor burden. The Stanford Institute of Medicine Animal Care and Use Committee approved all animal studies and procedures.

### Tumor dissociation, cell sorting, and freezing

Primary tumors and metastases were individually microdissected and dissociated using collagenase IV (Thermo Fisher), dispase (Corning), and trypsin (Invitrogen) at 37°C for 30 minutes. After dissociation, the samples remained on ice and in the presence of 2mM EDTA (Promega) and 1U/mL DNase (Sigma-Aldrich) to prevent aggregation. Cells were stained with antibodies to CD45 (30-F11), CD31 (390), F4/80 (BM8), and Ter119 (all from Biolegend) to exclude hematopoietic and endothelial linages (lineage-positive (Lin+) cells). DAPI was used to exclude dead cells. BD FACSAria sorters (BD Biosciences) were used for cell sorting. tdTomato+, Lin-, DAPI- cells were FACS sorted into microcentrifuge tubes, spun down at 500rcf for 5 minutes at 4°C in a fixed-angle centrifuge, re-suspended in 250uL freezing media (Bambanker; Wako Chemicals USA), and left at −80°C overnight before being transferred to liquid nitrogen storage until bulk ATAC-seq and scATAC-seq library preparation.

### ATAC-sequencing library preparation for primary tumors and metastases

FACS-isolated samples were taken out of storage in liquid nitrogen, quickly thawed at 37°C, diluted with 1mL PBS, and centrifuged at 300rcf for 5 minutes at 4°C in a fixed-angle centrifuge. Primary tumors and metastases were then processed for ATAC-seq library preparation using the same protocol used for cell lines, except the amount of transposase was decreased proportionally for samples with less than 50K cells. For example, for a sample with 10K cells, 1/5^th^ the normal amount of transposase was used in the 50uL transposition reaction. The remaining volume was replaced with ddH_2_O.

### RNA-sequencing library preparation for primary tumors and metastases

RNA was extracted from sorted cancer cells using the AllPrep DNA/RNA Micro Kit (Qiagen). RNA quality of each tumor sample was assessed using the RNA6000 PicoAssay for the Bioanalyzer 2100 (Agilent) as per the manufacturer’s instructions. All of the RNA used for RNA-seq had an RNA integrity number (RIN) > 8.0. RNA-sequencing libraries were generated as previously described ^41^ and sequenced using 200-cycle kits on an Illumina HiSeq 2000.

### scATAC-sequencing library preparation for primary tumors and metastases

FACS-isolated samples were taken out of storage in liquid nitrogen, quickly thawed at 37°C, diluted with 1mL PBS, and centrifuged at 300rcf for 5 minutes at 4°C in a fixed-angle centrifuge. Cells were re-suspended in PBS + 0.04% BSA, passed through a 40uM Flowmi cell strainer (Sigma) to minimize cell debris, and cell concentration was determined. Primary tumors and metastases were then processed for scATAC-seq library preparation according to standard droplet-based protocols (10x Genomics; Chromium Single Cell ATAC Library and Gel Bead Kit v1.0).

### scATAC-seq data processing and alignment

Raw sequencing data was converted to fastq format using cellranger atac mkfastq (10x Genomics, version 1.2.0). Single-cell ATAC-seq reads were aligned to the mm10 reference genome and quantified using cellranger count (10x Genomics, version 1.2.0). The current version of Cell Ranger can be accessed here: https://support.10xgenomics.com/single-cell-atac/software/downloads/latest.

### scATAC-seq data analysis – ArchR

We used ArchR^42^ for all downstream scATAC-seq analysis (https://greenleaflab.github.io/ArchR_Website/). We used the fragments files for each sample with their corresponding csv file with cell information. We then created Arrow files using “createArrowFiles” with using the barcodes from the sample 10x CSV file with “getValidBarcodes”. This step adds the accessible fragments a genome-wide 500-bp tile matrix and a gene-score matrix. We then added doublet scores for each single cell with “addDoubletScores” and then filtered with “filterDoublets”. Additionally, we then filtered cells that had a TSS enrichment below 6, less than 1,000 fragments or more than 50,000 fragments. We then reduced dimensionality with “addIterativeLSI” excluding chrX and chrY from this analysis. We then added clusters with “addClusters” with a resolution of 0.4. We then added a UMAP with “addUMAP” and minDist of 0.6. We identified 12 scATAC-seq clusters with this analysis. We then created a reproducible non-overlapping peak matrix with “addGroupCoverages” and “addReproduciblePeakSet”. We then quantified the number of Tn5 insertions per peak per cell using “addPeakMatrix”. We then added motif annotations using “addMotifAnnotations” with chromVAR mouse motifs version 1 “mouse_pwms_v1”. We then computed chromVAR deviations for each single cell with “addDeviationsMatrix”. For TF footprinting of NKX2-1 and SOX17 we used “plotFootprints” with normalization method “subtract” which substracts the Tn5 bias from the ATAC footprint.

To further characterize the 12 scATAC-seq clusters based on their metastatic state, we computed differential peaks from the LKB1-deficient bulk primary tumors and metastases. We took significantly differential peaks (|log2 Fold Change| > 3 and FDR < 0.01) and overlapped these peaks with the scATAC-seq peaks. The average accessibility and SEM across these overlapping regions was then plotted for peaks specific to primary tumors and peaks specific to metastases independently.

## Acknowledgments

We thank J. Sage, A. Trevino, and members of the Greenleaf and Winslow laboratories for helpful comments. We thank the Stanford Shared FACS facility, Veterinary Service Center, and P. Chu for technical support. We thank A. Orantes for administrative support.

## Author contributions

S.E.P., J.M.G., M.M.W., and W.J.G conceived the project and designed the experiments. S.E.P. led the experimental data production together with contribution from J.M.G., M.R.C., J.J.B., M.K.T., A.B.P., and R.T.. S.E.P. and J.M.G led the data analysis. S.E.P. performed the CRISPR screen analysis and RNA-seq analysis. J.M.G. and S.E.P. performed the ATAC-seq and scATAC-seq analysis. J.M.G was supervised by H.Y.C and W.J.G. S.E.P was supervised by M.C.B., W.J.G, and M.M.W.. S.E.P., J.M.G., W.J.G, and M.M.W. wrote the manuscript with input from all authors.

## Data and materials availability

Plasmids generated in this study are available from the Lead Contact without restriction. We have made available all raw sequencing, aligned fragments, and bigwigs for all ATAC-seq samples in **Supplementary Table 2** through AWS. Additionally, all matrices (peak matrix and chromVAR) are available in **Supplementary Table 2** through AWS. We also made the 10x cell ranger atac output files and all scATAC-seq matrices used in this study available in **Supplementary Table 2** through AWS. All sequencing data have been deposited in the Gene Expression Omnibus (GEO) as part of GEO accession GSE167381. To view please go to https://www.ncbi.nlm.nih.gov/geo/query/acc.cgi?acc=GSE167381 and enter token ubglmkeixbubzgj into the box. All gene expression matrices (count and tpm) are made available in **Supplementary Table 3** and **Supplementary Table 6**. The aligned CRISPR screen matrix is available in **Supplementary Table 1**.

## Code availability

All custom code used in this work is available upon request. We additionally will host a Github website that includes the main analysis code used in this study. This website will also include instructions for downloading and utilizing our supplemental matrices.

## Captions for Supplementary Tables 1-7

**Supplementary Table 1. (separate file)**

Enriched sgRNAs and gene targets in LKB1-restored cells compared to LKB1-deficient cells from genome-scale CRISPR/Cas9 screen.

**Supplementary Table 2. (separate file)**

List of all ATAC-seq and scATAC-seq samples processed in this study with related quality control information.

**Supplementary Table 3. (separate file)**

Gene expression changes in LKB1-restorable and LKB1-unrestorable cell lines after treatment with 4-OHT or vehicle.

**Supplementary Table 4. (separate file)**

sgRNA spacer sequences used in this study.

**Supplementary Table 5. (separate file)**

List of all *KPT* and *KPT;Lkb1^−/−^* mouse samples processed for ATAC-seq in this study.

**Supplementary Table 6. (separate file)**

Gene expression of LKB1-proficient *KPT* and LKB1-deficient *KPT;Lkb1^−/−^* mouse primary tumors and metastases.

**Supplementary Table 7. (separate file)**

Source data for panels throughout the text.

**Extended Data Figure 1.**
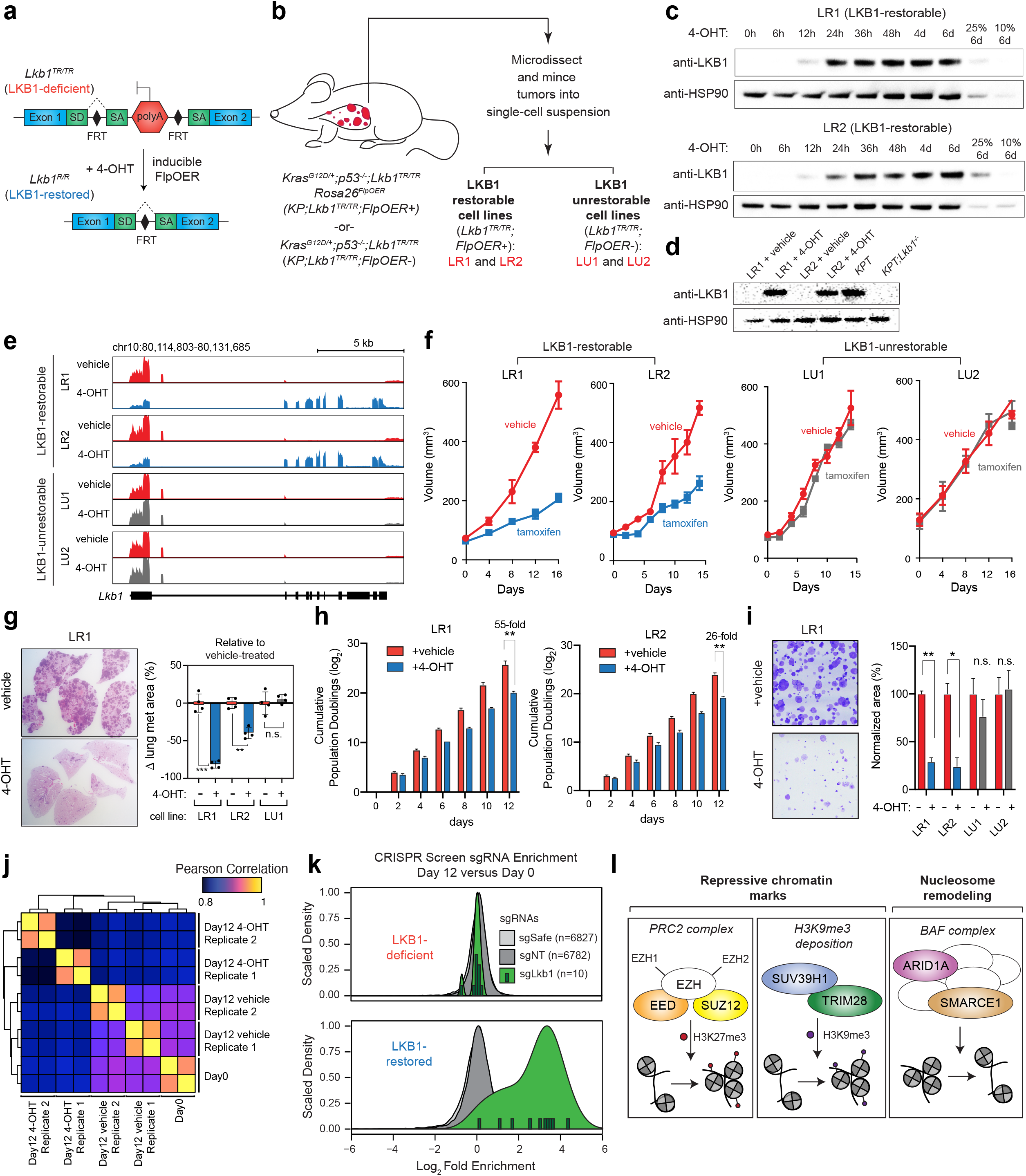
Validation and quality control of inducible LKB1 restoration model and genome-scale CRISPR/Cas9 screen. **a.** Schematic of the homozygous, restorable *Lkb1^TR/TR^* alleles in lung adenocarcinoma cell lines. SA = splice acceptor, SD = splice donor, FRT = flippase recognition target. **b.** Schematic of the derivation of LKB1-restorable *Lkb1^TR/TR^;FlpOER+* cell lines (LR1 and LR2) and LKB1-unrestorable *Lkb1^TR/TR^;FlpO-ER-* cell lines (LU1 and LU2). **c.** Expression of LKB1 by immunoblot over a time-course of 4-OHT treatment in LR1 cells (*top*) and LR2 cells (*bottom*), represented in hours (h) and days (d). HSP90 is a sample processing control. 25% and 10% of input after six days of 4-OHT treatment is shown for a visual comparison. **d.** Expression of LKB1 by immunoblot in LR1 and LR2 cells treated with vehicle or 4-OHT for six days compared to a *KPT* cell line and a *KPT;Lkb1^−/−^* cell line. HSP90 is a sample processing control. **e.** RNA-sequencing reads mapping to the *Lkb1* locus for the indicated cell lines following six days of 4-OHT or vehicle treatment. **f.** Subcutaneous growth assay following injection of the indicated cell lines into recipient NSG mice. Tamoxifen or vehicle treatment was initiated on day 0. Tamoxifen is metabolized in the liver following oral gavage administration to form 4-OHT. Mean tumor volume as measured by calipers of six tumors per condition +/− SEM is shown. **g.** Intravenous (i.v.) transplant assays of the indicated cell lines treated with 4-OHT or vehicle for six days, injected i.v. into recipient NSG mice, and analyzed after 21 days. *Left*: Representative lung histology 21 days after i.v. transplantation of LR1 cells. *Right*: Change in % tumor area as quantified by histology in each cell line compared to the same cell line treated with vehicle. Mean area of four mice per condition +/− SEM is shown. **p < 0.001, ***p < 0.0001, n.s. = not significant. **h.** Cumulative population doublings in log-scale recorded over 12 days for the LR1 and LR2 cell lines treated with 4-OHT or vehicle for 12 days. Each cell line and treatment group was cultured and analyzed in triplicate. Mean +/− SEM is shown. All comparisons between 4-OHT-treated and vehicle-treated cells beyond two days were significant (p < 0.001) for LR1. All comparisons between4-OHT-treated and vehicle-treated cells beyond two days (p < 0.01) and four days (p < 0.001) were significant in LR2. **i.** Clonogenic growth assays in the indicated cell lines treated with 4-OHT or vehicle for six days. *Left*: Representative image of clonogenic growth in LR1 cells. *Right*: % normalized area of cell growth. Each treatment group was cultured and analyzed in triplicate. Mean +/− SEM is shown. *p < 0.01, **p < 0.001, n.s. = not significant. **j.** Heatmap of Pearson correlation matrix of log-normalized counts across all samples in the genome-scale CRISPR/Cas9 screen. **k.** Log_2_ fold enrichment of 13,609 negative control sgRNAs (sgSafe in light grey and sgNT in dark grey) and 10 positive control sgRNAs targeting *Lkb1* (sgLkb1 in light green, individual sgRNAs in dark green) at day 12 versus day 0 of the genome-scale CRISPR/Cas9 screen in two vehicle-treated, LKB1-deficient replicates (*top*) and two 4-OHT-treated, LKB1-restored replicates (*bottom*). **i.** Schematic outlining the functions of the top six chromatin modifiers identified in the screen: EED, SUZ12, SUV39H1, TRIM28, ARID1A, and SMARCE1 (FDR < 0.1).

**Extended Data Figure 2.**
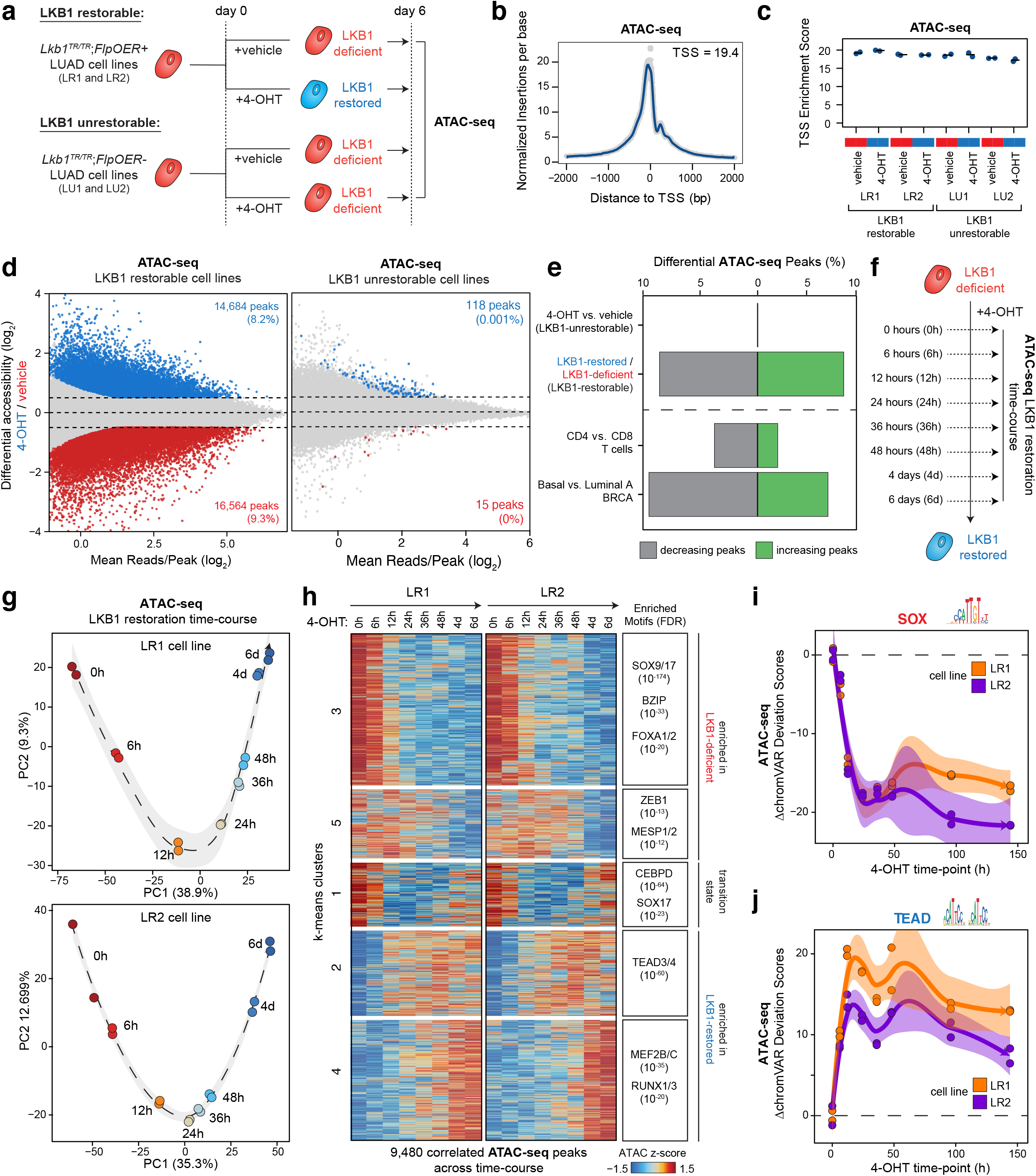
LKB1 restoration drives widespread changes in chromatin accessibility in lung adenocarcinoma cells. **a.** Schematic of preparing LKB1-deficient and LKB1-restored samples prior to ATAC-seq library preparation. Cell lines were treated with 4-OHT or vehicle for six days. **b.** Representative plot of aggregate signal around the transcription start site (TSS) for all ATAC-seq peaks in one vehicle-treated, LR1 replicate. This plot represents the signal-to-noise quantification of our ATAC-seq data. TSS enrichment scores greater than 10 indicate high quality ATAC-seq data. **c.** TSS enrichment scores for 16 ATAC-seq libraries with technical replicates. **d.** Differential accessibility across 178,783 ATAC-seq peaks following 4-OHT treatment in the LKB1-restorable (LR1 and LR2) and LKB1-unrestorable (LU1 and LU2) cell lines. The x-axis represents the log_2_ mean accessibility per peak and the y-axis represents the log_2_ fold change in accessibility following 4-OHT treatment. Colored points are significant (|log_2_ fold change|>0.5, FDR <0.05). **e.** Percentage of differential peaks (|log_2_ fold change|>0.5, FDR <0.05) across multiple ATAC-seq comparisons. **f.** Schematic of preparing samples for LKB1-restoration time-course. Cell lines were treated with 4-OHT for eight different time-points (0 hours, 6 hours, 12 hours, 24 hours, 36 hours, 48 hours, 4 days, and 6 days) in two cell lines (LR1 and LR2). **g** and **h.** PCA (**g**) and k-means clustering (**h**) of 9,480 correlated, variable ATAC-seq peaks across the LKB1 restoration time-course in two cell lines (LR1 and LR2). Each row of the heatmap represents a z-score of log_2_ normalized accessibility across all samples within each cell line. **i** and **j.** SOX (**i**) and TEAD (**j**) motif accessibility changes (ΔchromVAR deviation scores) across time in two cell lines (LR1 and LR2) treated with 4-OHT for the indicated time-points.

**Extended Data Figure 3.**
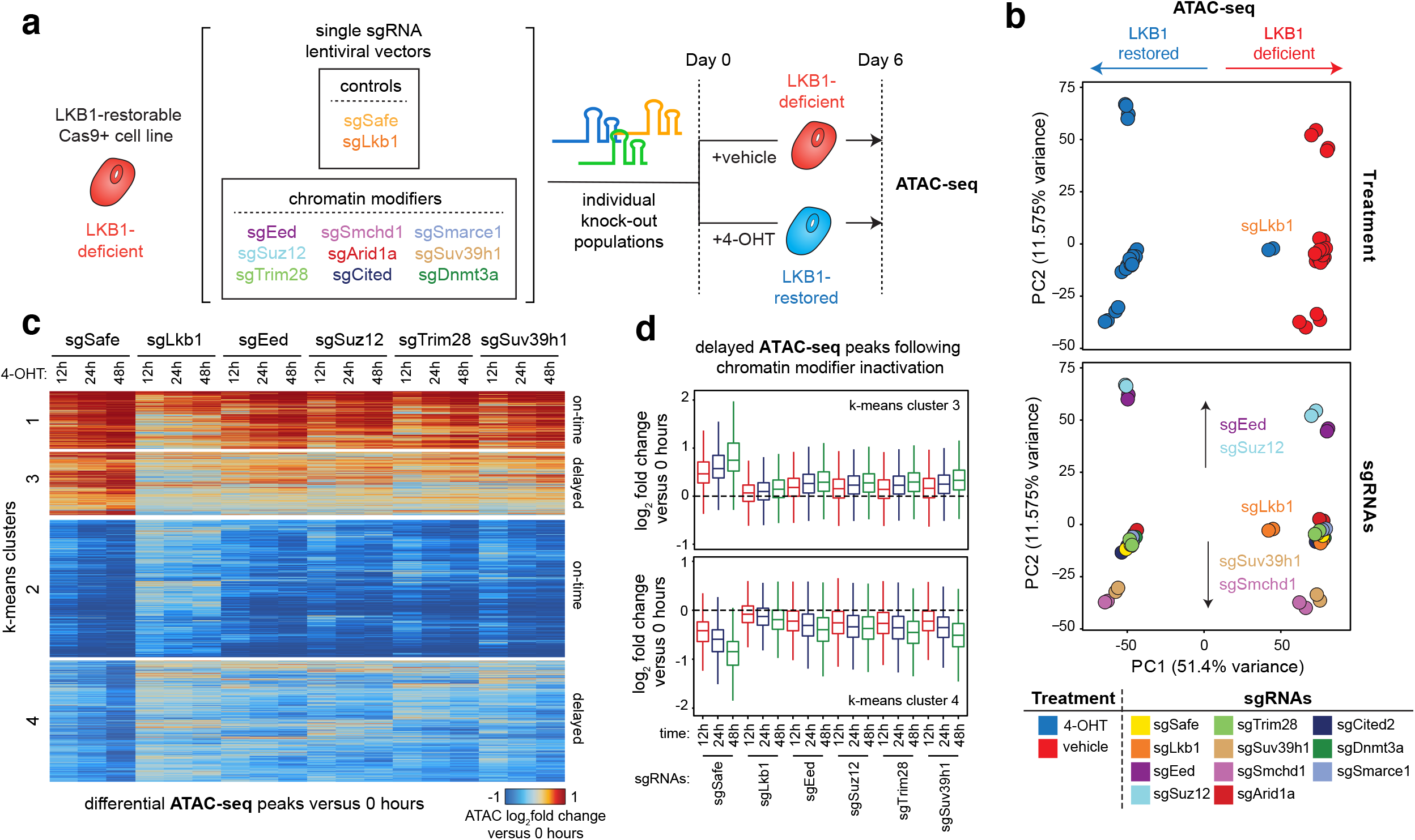
Inactivating chromatin modifiers only delays LKB1-induced chromatin changes. **a.** Schematic of generating single knock-out populations of chromatin modifiers identified in the CRISPR screen, treating with 4-OHT or vehicle for six days, and processing for ATAC-seq. **b.** Principle component analysis (PCA) of the top 10,000 variable ATAC-seq peaks across the indicated LR1;Cas9 knock-out populations treated with 4-OHT or vehicle. **c.** K-means clustered heatmap of differential peak accessibility (log_2_ fold change) for each genotype of LR1;Cas9 cells treated with 4-OHT for up to 48 hours compared to 0 hours. All peaks differential between sgSafe (0 hours 4-OHT) and sgSafe (48 hours 4-OHT) are shown. Each row represents the log_2_ fold change of each genotype and time-point versus the same genotype’s initial time-point (day 0). **d.** Log_2_ fold change in mean peak accessibility for all peaks in k-means cluster 3 (top) and cluster 4 (bottom) from (**c**) for the indicated genotype and 4-OHT time-points compared to 0 hours 4-OHT.

**Extended Data Figure 4.**
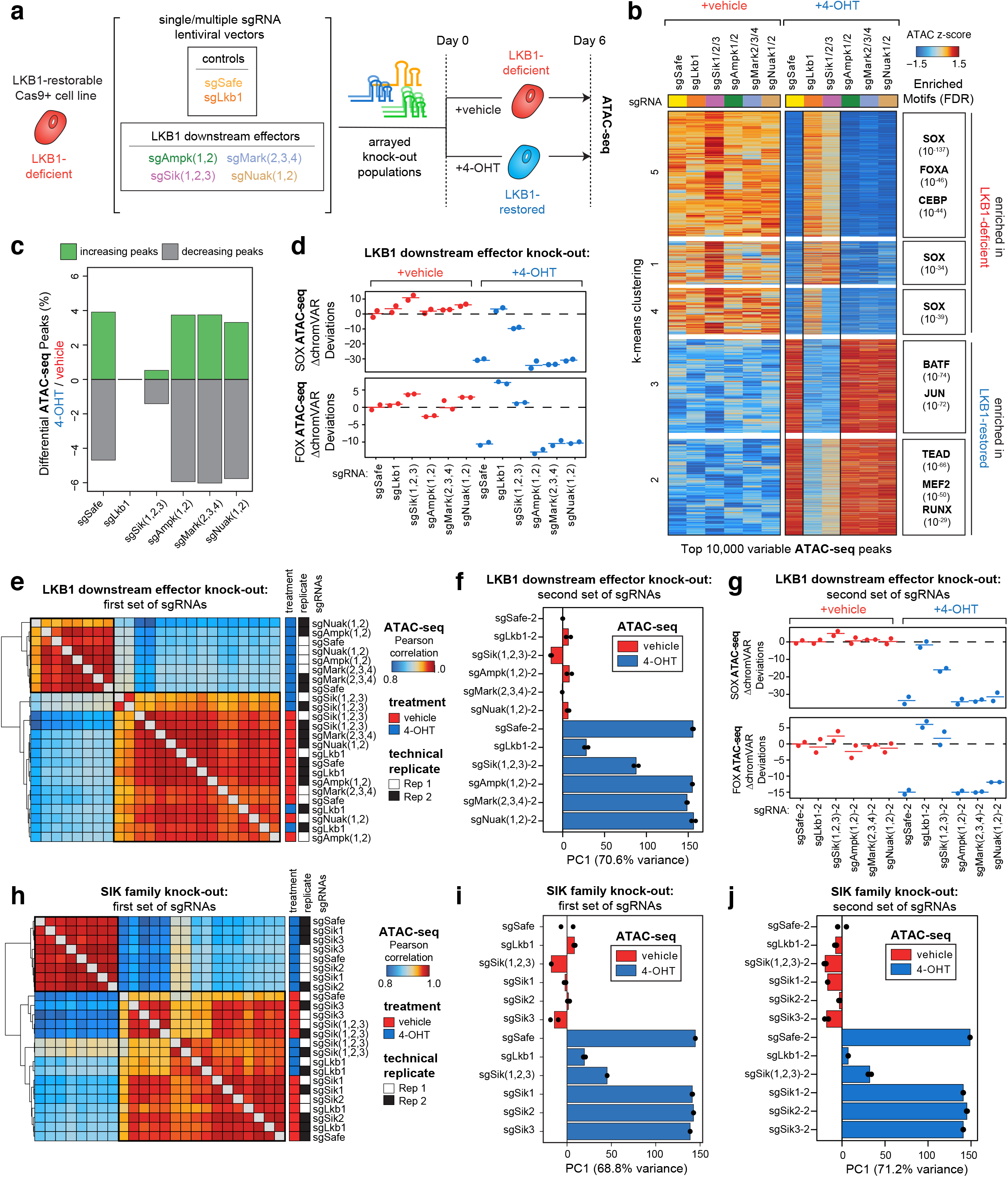
SIK family members mediate LKB1-induced chromatin changes. **a.** Schematic of generating single and multiple sgRNA knock-out cell lines and processing for ATAC-seq. LR1;Cas9 cells were treated with 4-OHT or vehicle for six days. **b.** *Left*: Heatmap of peak accessibility between each knock-out population treated with 4-OHT or vehicle. Each row represents a z-score of log_2_ normalized accessibility across all samples. *Right*: Transcription factor hypergeometric motif enrichment in each k-means cluster. **c.** Percent of differential ATAC-seq peaks (|log_2_ fold change|>0.5, FDR <0.05) across LKB1-restorable cells treated with 4-OHT or vehicle. **d.** SOX (*top*) and FOXA (*bottom*) motif accessibility changes (ΔchromVAR deviation scores normalized to vehicle-treated sgSafe) across LKB1-restorable knock-out populations treated with 4-OHT or vehicle. **e.** Heatmap of Pearson correlation matrix of log_2_-normalized accessibility (in counts per million (CPM)) across LKB1 downstream effector knock-out genotypes with and without LKB1 restoration in LR1;Cas9 cells. **f.** PCA of the top 10,000 variable ATAC-seq peaks across LR1;Cas9 knock-out populations treated with 4-OHT or vehicle. Principle components besides PC1 (70.6%) account for <4% of the variance in the dataset. **g.** SOX (*top*) and FOXA (*bottom*) motif accessibility changes (ΔchromVAR deviation scores normalized to vehicle-treated sgSafe) across LKB1-restorable knock-out populations treated with 4-OHT or vehicle. **h.** Heatmap of Pearson correlation matrix of log_2_-normalized accessibility (in counts per million (CPM)) across LKB1 downstream effector knock-out genotypes with and without LKB1 restoration in LR1;Cas9 cells. **i** and **j.** PCA of the top 10,000 variable ATAC-seq peaks across LR1;Cas9 knock-out populations treated with 4-OHT or vehicle. Principle components besides PC1 account for <4% of the variance in the dataset.

**Extended Data Figure 5.**
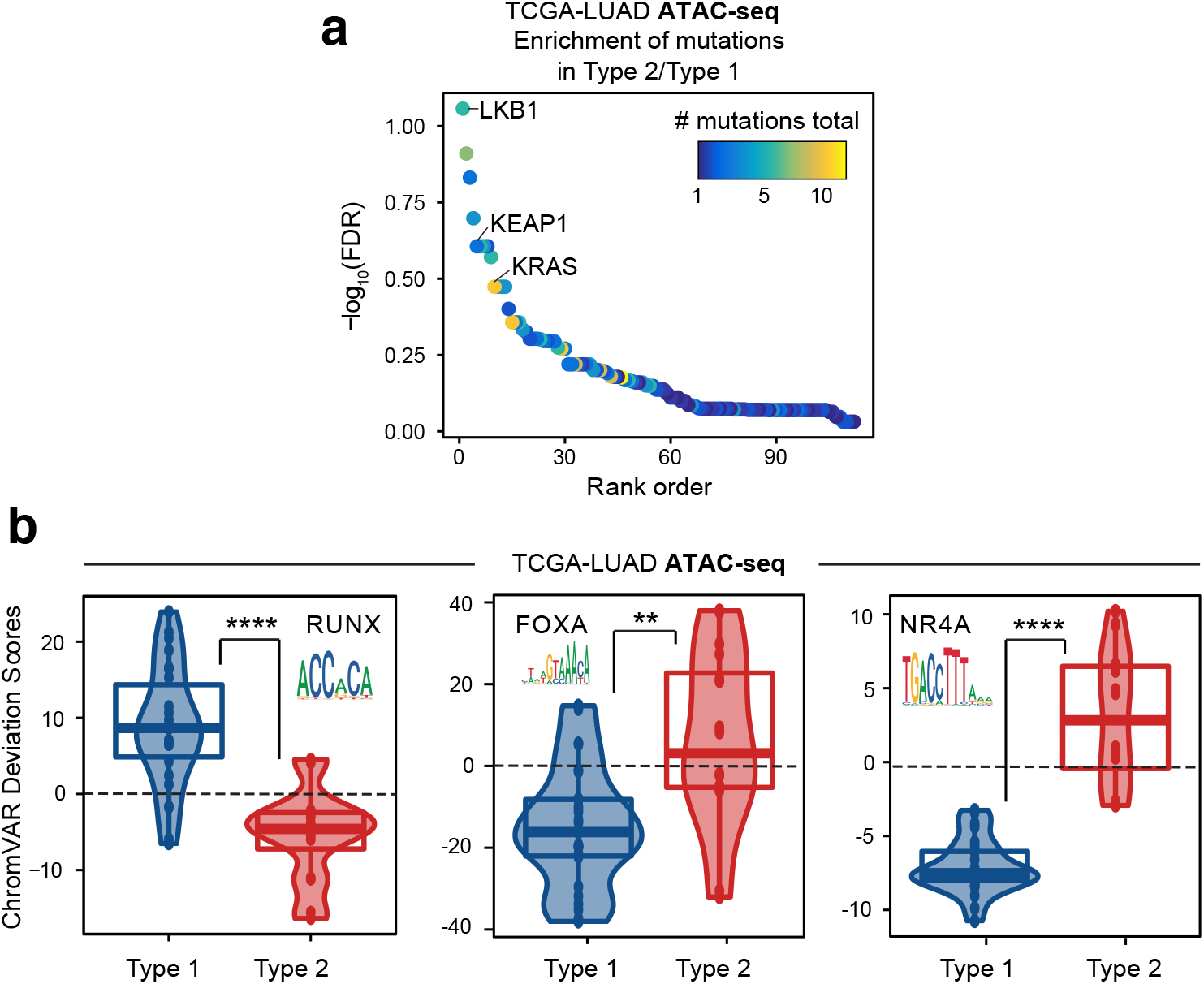
Loss of LKB1 partitions human lung adenocarcinoma primary tumors into two chromatin sub-types. **a.** Enrichment of mutations in Chromatin Type 2 tumors compared to Chromatin Type 1 tumors. Genes are ranked according to −log_10_(FDR), with Rank 1 (LKB1) being the most significant (see Methods), as indicated on the y-axis. Points are colored by the number of mutations in the TCGA-LUAD ATAC-seq dataset (out of 21 samples). **b.** ChromVAR deviation scores for the indicated transcription factor motifs for samples in the TCGA-LUAD ATAC-seq dataset. *p < 0.1, **p < 0.005, ****p< 10^−6^ using a two-sided t-test.

**Extended Data Figure 6.**
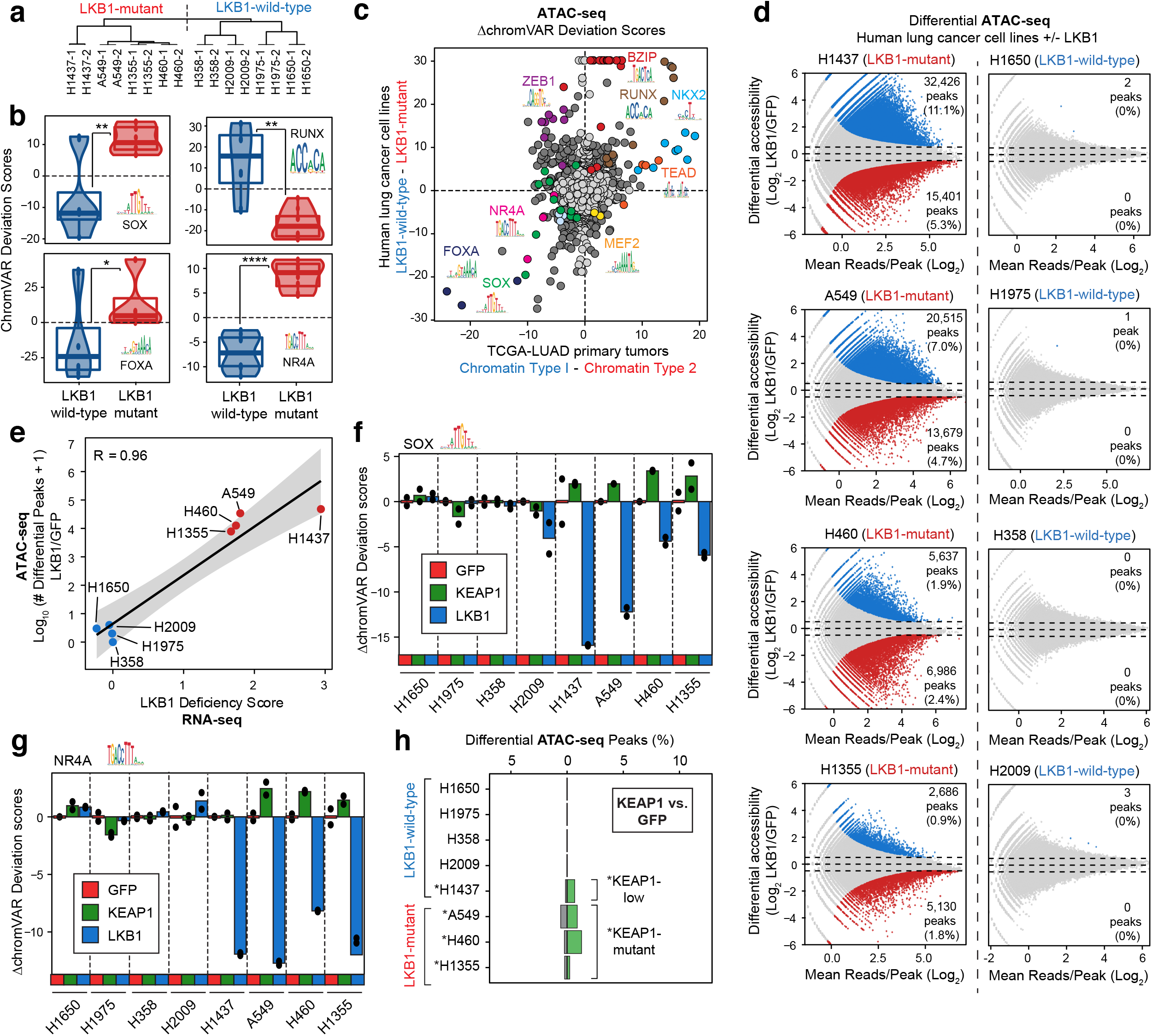
Loss of LKB1 drives a unique chromatin state in human lung adenocarcinoma cell lines. **a.** Hierarchical clustering of human lung cancer cell lines using the Euclidian distance within the first three principle components from Fig. 2d. **b.** ChromVAR deviation scores for the indicated transcription factor motifs in eight human lung cancer cell lines at baseline. *p<0.1,**p<0.005, ****p<10^−6^ using a two-sided t-test. **c.** Comparison of the changes in motif accessibility (Δ chromVAR deviation scores) across LKB1-wild-type and LKB1-mutant human lung cancer cell lines (y-axis) and Chromatin Type 1 and Type 2 tumors (x-axis). Dark grey or colored points are called significantly different (q < 0.05) across both comparisons. Light grey points are not significant. A selection of motif families and their associated motif logos are indicated. **d.** Differential accessibility across ATAC-seq peaks following LKB1 wild-type expression in eight human lung cancer cell lines. The x-axis represents the log_2_ fold change in accessibility following LKB1 restoration. LKB1-mutant and LKB1-wild-type status at baseline is indicated. Colored points are significant (|log_2_ fold change|>0.5, FDR <0.05). **e.** LKB1-deficiency score by RNA-seq (using 16-gene signature from Kaufmann et al., 2017) compared to log_10_(number of differential ATAC-seq peaks + 1) following LKB1 expression in each indicated cell line. Pearson correlation indicated in top left. **f** and **g.** Relative chromVAR deviation scores for SOX (**f**) an NR4A (**g**) motifs in the indicated cell lines transduced with GFP, LKB1, or KEAP1. Scores are normalized based on the GFP control for each cell line. **h.** Percent of differential ATAC-seq peaks (|log_2_ fold change|>0.5, FDR <0.05) in cells transduced to express KEAP1 compared to GFP.

**Extended Data Figure 7.**
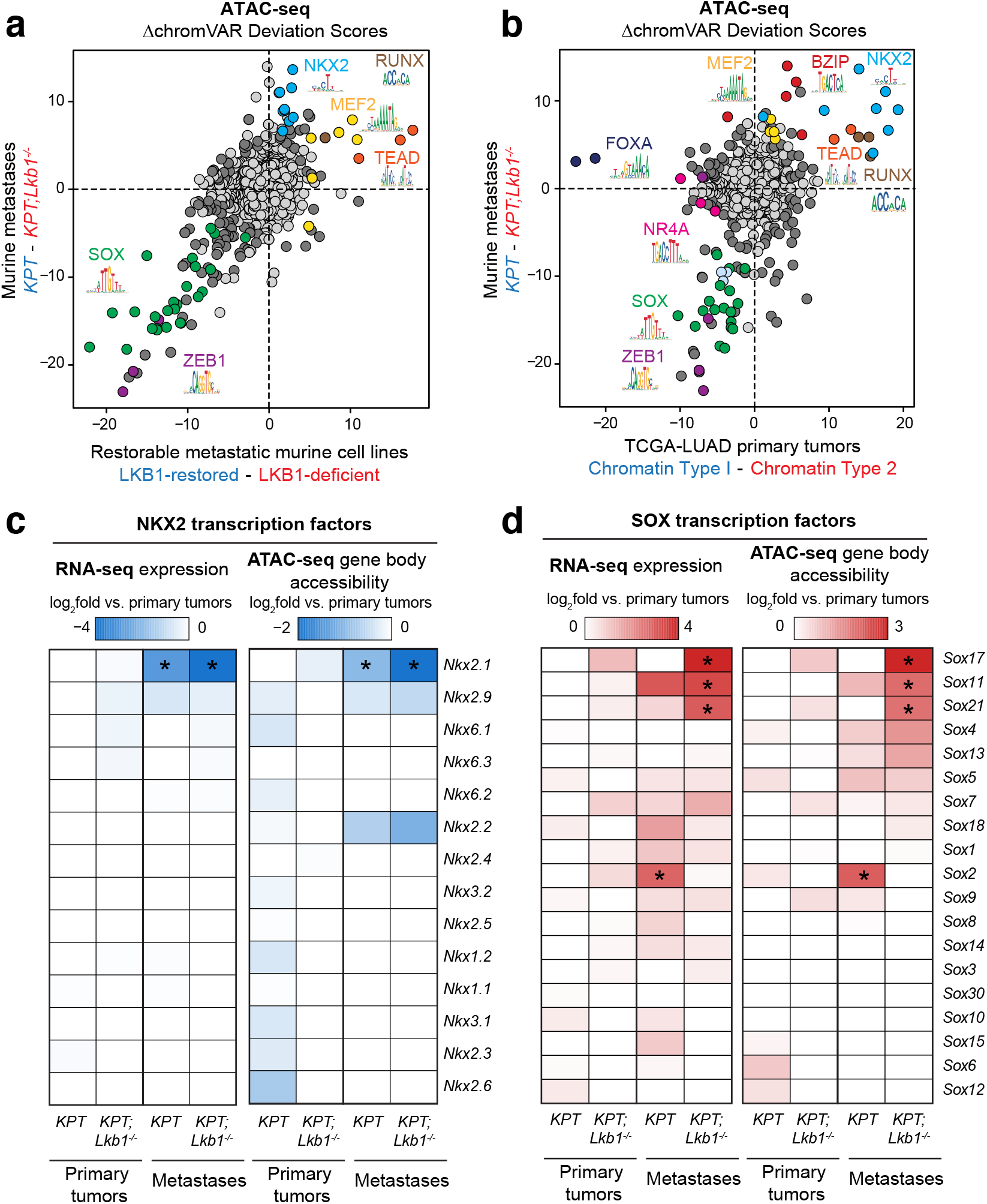
Genotype-specific activation of SOX17 in LKB1-deficient metastatic cells. **a** and **b.** Comparison of the changes in motif accessibility (ΔchromVAR deviation scores) between murine LKB1-proficient (*KPT*) and LKB1-deficient (*KPT;Lkb1^−/−^*) metastases (y-axis) and between murine LKB1-restored and LKB1-deficient cells (x-axis; **b**) or Chromatin Type 1 tumors and Chromatin Type 2 tumors(x-axis; **c**). Dark grey or colored points are called significantly different (q < 0.05) across both comparisons. Light grey points are not significant. A selection of motif families and their associated motif logos are indicated. **c.** log_2_ fold change in mRNA expression (*left*) and accessibility within the gene body (*right*) of each NKX2 transcription factor compared to the average expression and accessibility in primary tumor samples. Asterisks indicate transcription factors with greater than log_2_fold change of −1 in both RNA and ATAC measurements. **d.** log_2_ fold change in mRNA expression (*left*) and accessibility within the gene body (*right*) of each SOX transcription factor compared to the average expression and accessibility in primary tumor samples. Asterisks indicate transcription factors with greater than log_2_fold change of 2 in both RNA and ATAC measurements.

**Extended Data Figure 8.**
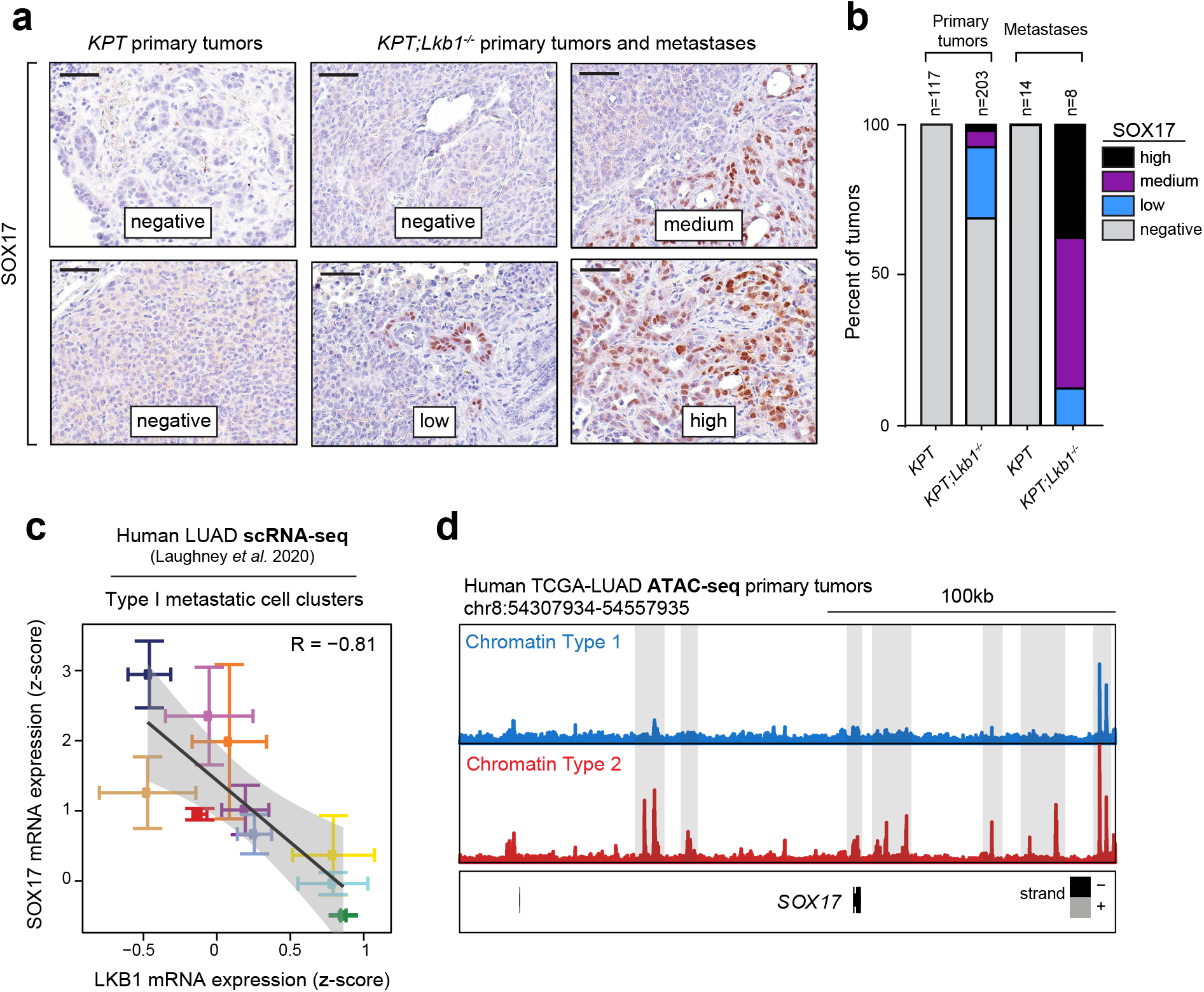
LKB1-deficient primary tumors harbor sub-populations of SOX17+ cells. **a.** Representative immunohistochemistry (IHC) against SOX17 and grading of SOX17 expression for LKB1-proficient *KPT* and LKB1-deficient *KPT;Lkb1^−/−^* samples. Images are annotated according to percent area of the tumor composed of SOX17+ cells. Negative (0%), low (<25%), medium (25-50%), and high (>50%). Scale bars represent 50uM. **b.** Quantitation of SOX17 protein expression in LKB1-proficient *KPT* and LKB1-deficient *KPT;Lkb1^−/−^* primary tumors and metastases, graded according to (**a**). The number of samples analyzed for histology for each genotype and tumor type is indicated at the top. Overall 0% of LKB1-proficient primary tumors or metastases had SOX17+ cells, 31% of LKB1-deficient primary tumors had SOX17+ cells, and 100% of LKB1-deficient metastases had SOX17+ cells. **c.** Correlation of *SOX17* mRNA expression (y-axis) and *LKB1* mRNA expression (x-axis) in ten human lung adenocarcinoma samples that contain Type 1 metastatic cell clusters (H0 and H3; Laughney et al. 2020). Each point indicates the mean value of *SOX17* or *LKB1* expression for each sample +/− SEM for all single cells evaluated by scRNA-seq. **d.** *SOX17* genome accessibility track of the average ATAC-seq signal from Chromatin Type 1 and Chromatin Type 2 tumors.

**Extended Data Figure 9.**
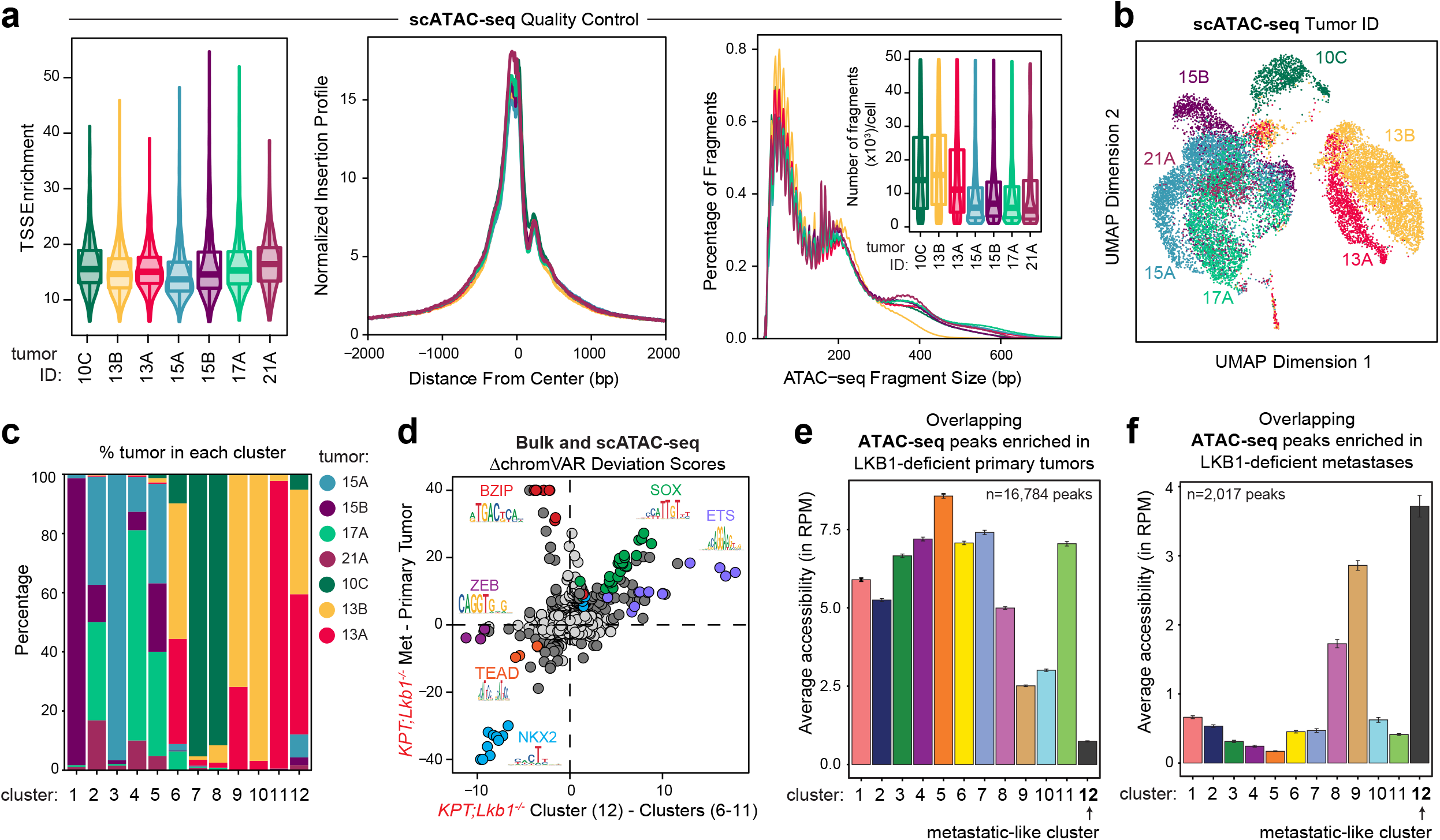
A subset of LKB1-deficient primary tumors harbor metastatic-like, SOX17+ sub-populations. **a.** scATAC-seq quality control metrics. TSS enrichment (*left, middle*), insertion profiles (*right*), and number of fragments per cell (*right inset*) in each of the seven primary tumors evaluated. **b.** Uniform Manifold Approximation and Project (UMAP) of cells from seven primary tumors, colored by tumor of origin. 10C, 13B, and 13A originated from three separate LKB1-deficient *KPT;Lkb1^−/−^* primary tumors. 17A, 15A, 21A, and 15B originated from four separate LKB1-proficient *KPT* primary tumors. **c.** Percent of cells from each cluster contained within each individual tumor sample. **d.** Comparison of the changes in motif accessibility (ΔchromVAR deviation scores) between LKB1-deficient *KPT;Lkb1^−/−^* metastases and primary tumors (y-axis) versus the average difference between cluster 12 cells and cells in clusters 1-11 (x-axis). Dark grey or colored points are called significantly different (q<0.05) across both comparisons. Light grey points are not significant. A selection of motif families and their associated motif logos are indicated. **e** and **f.** Average accessibility (in reads per million (RPM)) of the peaks in each scATAC-seq cluster that are enriched in LKB1-deficient primary tumors compared to LKB1-deficient metastases (**e**) or enriched in LKB1-deficient metastases compared to LKB1-deficient primary tumors (**f**) and are overlapping with the scATAC-seq peakset. Error bars indicate +/− SEM for each cluster’s average accessibility.

**Extended Data Figure 10.**
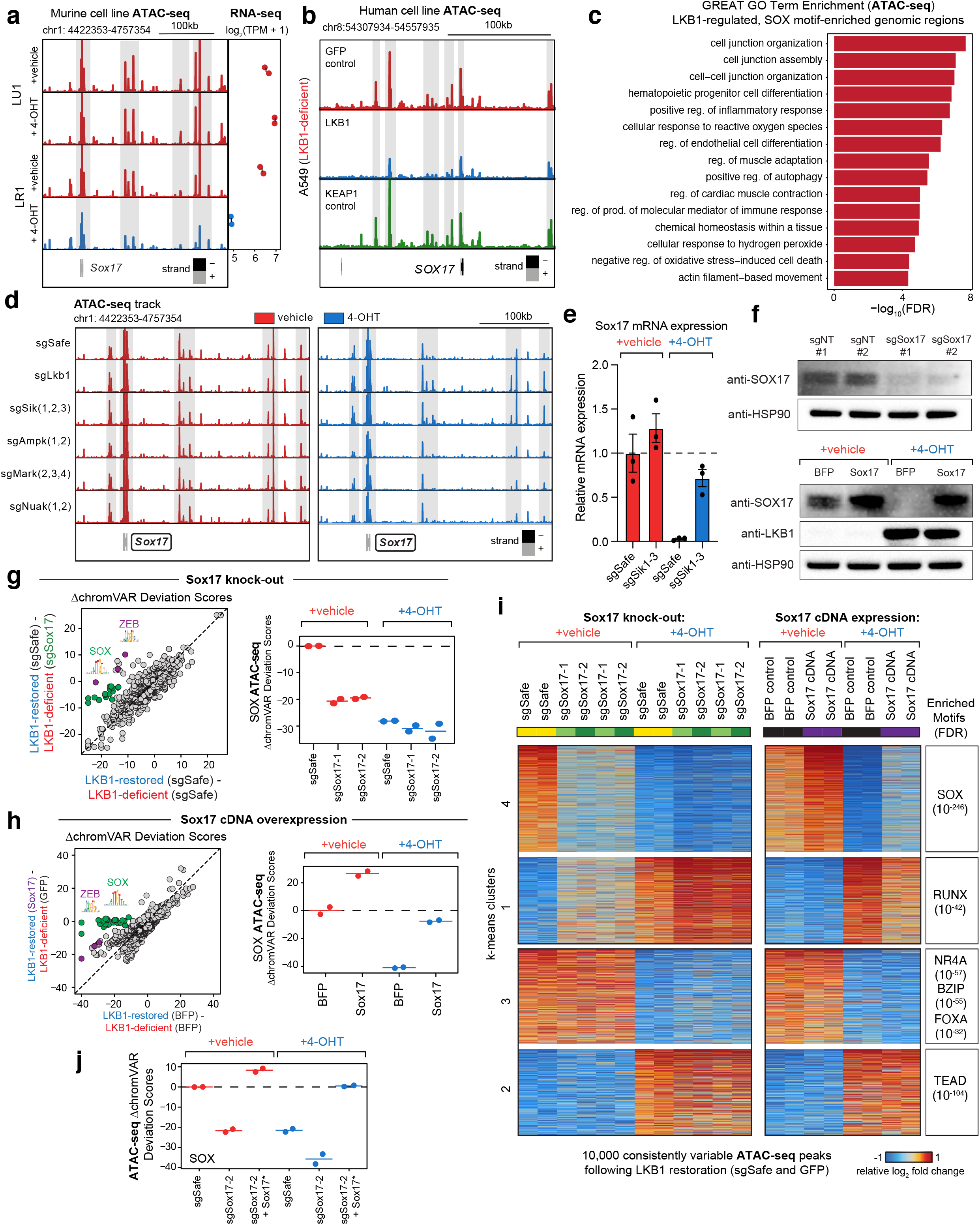
SOX17 regulates chromatin accessibility state in metastatic, LKB1-deficient cells. **a.** *Sox17* genome accessibility track (*left*) and mean *Sox17* mRNA expression (*right*) of an LKB1-unrestorable cell line (LU1) and an LKB1-restorable cell line (LR1) treated with 4-OHT or vehicle for six days. Highlighted in grey are significantly differential ATAC-seq peaks (log_2_ fold change < −0.5, FDR < 0.05) following LKB1 restoration. *Sox17* also has significantly decreased mRNA expression (log_2_ fold change < −1, FDR < 0.05) following LKB1 restoration. **b.** *SOX17* genome accessibility track of an LKB1-deficient cell line (A549) transduced with GFP, LKB1, or KEAP1. **c.** GREAT GO term enrichment of genes nearby the differential peaks that contain SOX binding motifs that are enriched in LKB1-deficient cells compared to LKB1-restored cells. **d.** *Sox17* genome accessibility track of an LKB1-restorable cell line (LR1;Cas9) transduced with lentiviral constructs containing sgSafe, sgSik1-3, sgAmpk1/2, sgMark2-4, or sgNuak1/2 and treated with 4-OHT or vehicle for six days. **e.** Relative mRNA expression of Sox17 in LR1;Cas9 cells transduced with lentiviral constructs containing sgSafe or sgSik1-3 and treated with either vehicle or 4-OHT for six days. **f.** Expression of of SOX17 and/or LKB1 by immunoblot in LR2;Cas9 cells transduced with non-targeting (sgNT#1 and sgNT#2) or Sox17-targeting sgRNAs (sgSox17#1 and sgSox17#2) (top) or LR2;Cas9 cells transduced with BFP-overexpressing (control) or Sox17-overexpressing constructs and treated with vehicle or 4-OHT for six days. HSP90 is a sample processing control. **g.** *Left:* Comparison of the changes in motif accessibility (ΔchromVAR deviation scores) between LKB1-restored and LKB1-deficient LR2;Cas9 cells transduced with sgSafe (x-axis) compared to the average differences after knocking out Sox17 in LKB1-deficient cells (y-axis). Dark grey or colored points are called significantly different (q<0.05) across both comparisons. *Right*: ChromVAR deviation scores for SOX motifs in each group (normalized to vehicle-treated sgSafe). Each point represents an ATAC-seq technical replicate, bar represents the mean. **h.** *Left:* Comparison of the changes in motif accessibility (ΔchromVAR deviation scores) between LKB1-restored and LKB1-deficient LR2;Cas9 cells transduced with a blue fluorescent protein (BFP) control (x-axis) versus the difference after overexpressing *Sox17* in LKB1-restored cells (y-axis). Dark grey or colored points are called significantly different (q<0.05) across both comparisons. *Right*: ChromVAR deviation scores for SOX motifs in each group (normalized to vehicle-treated BFP). Each point represents an ATAC-seq technical replicate, bar represents the mean. **i.** Heatmap of the relative log_2_fold changes of the indicated genotypes of cells with and without LKB1 restoration in LR2;Cas9 cells. The top 10,000 consistent, variable ATAC-seq peaks following LKB1 restoration in both sgSafe and BFP transduced cells are shown. Clusters 3 and 4 from the *Sox17* knock-out experiment are shown independently for emphasis in Fig. 5c. **j.** ChromVAR deviation scores for SOX motifs in each group in another cell line (LR1;Cas9). Each individual point represents an ATAC-seq technical replicate, bar represents the mean. sgSox17-2 + Sox17* indicates that the cells were transduced with a construct containing a sgRNA targeting Sox17 as well as a Sox17 cDNA that is resistant to sgRNA cutting (see Methods).

**Extended Data Figure 11.**
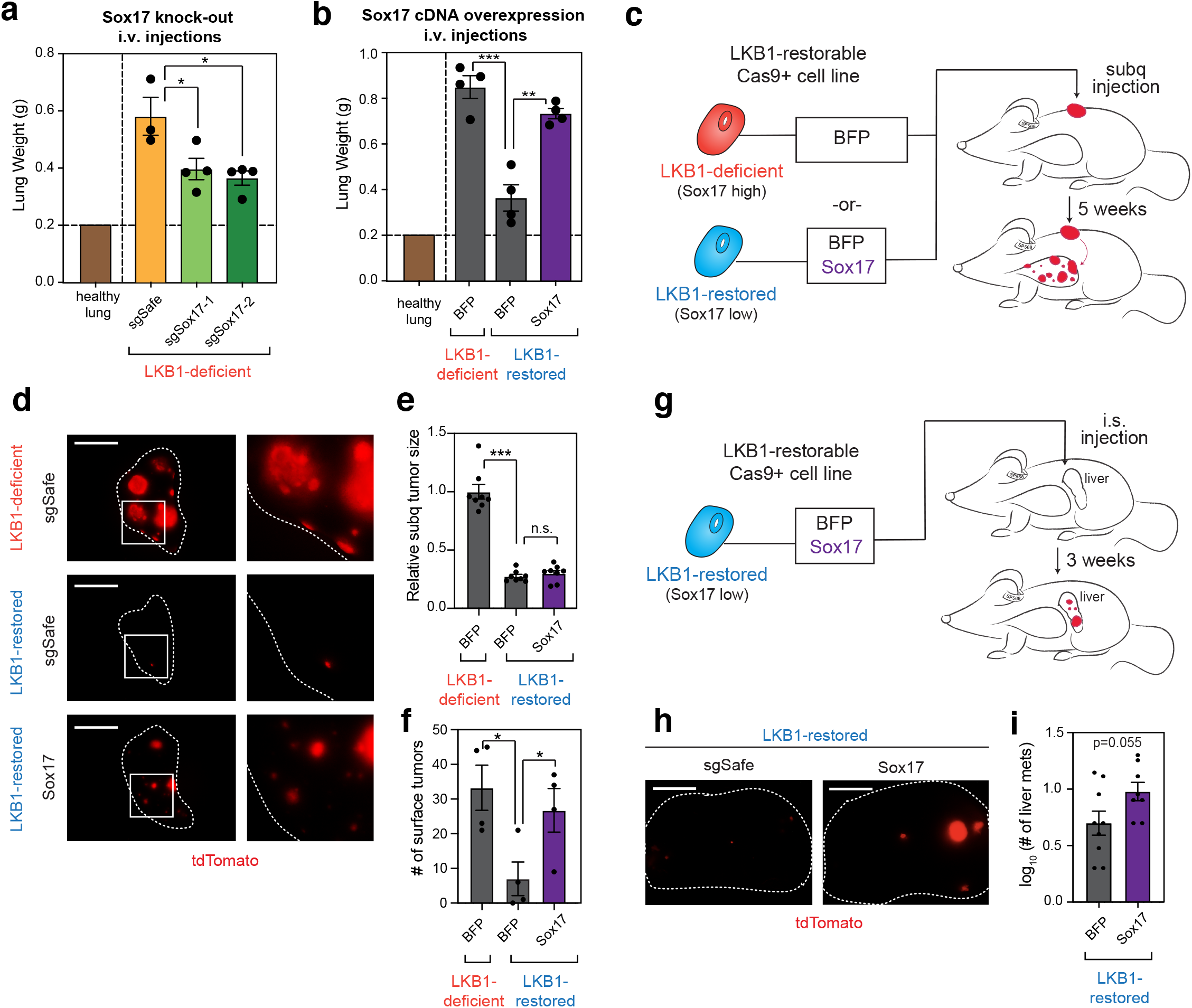
SOX17 regulates growth and chromatin state in metastatic, LKB1-deficient cells. **a** and **b.** Lung weight following injection of LR2;Cas9 cells treated with either vehicle (LKB1-deficient) or 4-OHT (LKB1-restored) after *Sox17* knock-out (**a**) or *Sox17* overexpression (**b**). *p<0.05, **p<0.005, ***p<0.0005. **c.** Schematic of injecting LKB1-deficient cells (LR2) expressing BFP or injecting LKB1-restored cells (LR2) expressing Sox17 cDNA or BFP subcutaneously (subq) into immunocompromised NSG mice. Metastatic tumor burden to the lung was analyzed five weeks post-injection. **d.** Representative fluorescent tdTomato+ images of single lung lobes following subq injection as outlined in (**c**). **e.** Relative tumor size following subcutaneous injection of the indicated cells. Each point represents an individual tumor and two tumors were injected per mouse. Condition +/− SEM is shown. **** p < 0.0005, n.s. = not significant. **f.** Number of surface tumors observed in the five lung lobes following subcutaneous injection of the indicated cells. Each point represents tumors evaluated from an individual mouse. Condition +/− SEM is shown. *p < 0.05. **g.** Schematic of injecting LKB1-deficient cells expressing BFP or injecting LKB1-restored cells expressing Sox17 cDNA or BFP intrasplenically (i.s.) into immunocompromised NSG mice. Metastatic tumor burden to the liver was analyzed three weeks post-injection. **h.** Representative fluorescent tdTomato+ images of the left lateral lobe of the liver following intrasplenic injection as outlined in (**g**). **i.** Log_10_ (number of liver metastases) following intrasplenic injection of the indicated cells. Each point represents tumors evaluated from an individual mouse. Condition +/− SEM is shown.

**Extended Data Figure 12.**
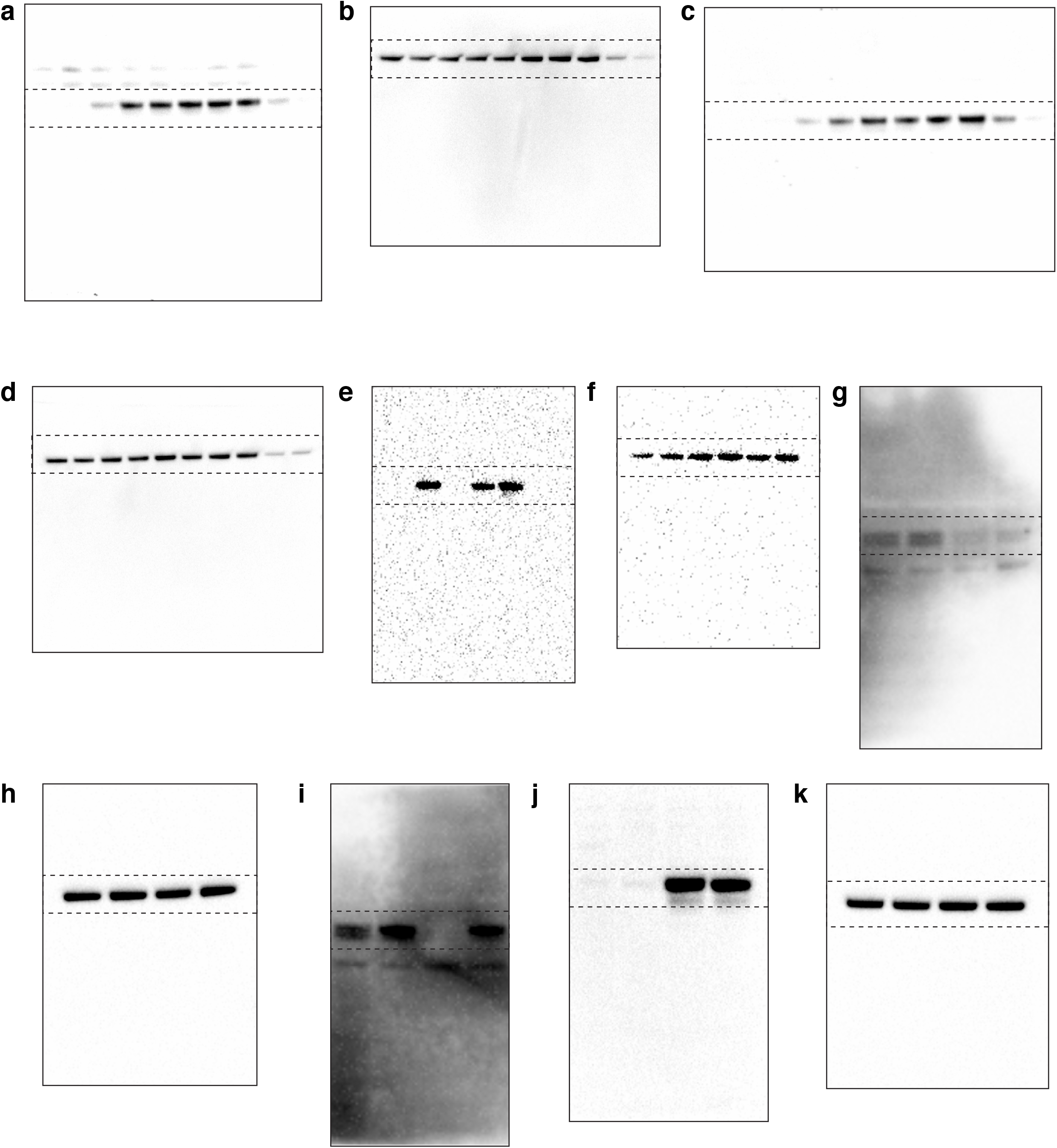
Source data for immunoblot analyses used in the text. **a.** Anti-LKB1 in Extended Data Figure 1c (LR1). **b.** Anti-HSP90 in Extended Data Figure 1c (LR1). HSP90 is a sample processing control. **c.** Anti-LKB1 in Extended Data Figure 1c (LR2). **d.** Anti-HSP90 in Extended Data Figure 1c (LR2). HSP90 is a sample processing control. **e.** Anti-LKB1 in Extended Data Figure 1d.**f.** Anti-HSP90 in Extended Data Figure 1d. HSP90 is a sample processing control. **g.** Anti-SOX17 in Extended Data Figure 10f (top). **h.** Anti-HSP90 in Extended Data Figure 10f (top). HSP90 is a sample processing control. **i.** Anti-SOX17 in Extended Data Figure 10f (bottom). **j.** Anti-LKB1 in Extended Data Figure 10f (bottom). **k.** Anti-HSP90 in Extended Data Figure 10f (bottom). HSP90 is a sample processing control.

